# Dynamic and structural insights into allosteric regulation on MKP5 a dual-specificity phosphatase

**DOI:** 10.1101/2024.09.05.611520

**Authors:** Erin Skeens, Federica Maschietto, Ramu Manjula, Shanelle Shillingford, Elias J. Lolis, Victor S. Batista, Anton M. Bennett, George P. Lisi

**Affiliations:** Department of Molecular Biology, Cell Biology and Biochemistry, Brown University, Providence, Rhode Island, USA; Department of Chemistry, Yale University, New Haven, Connecticut, USA; Department of Pharmacology, Yale School of Medicine, Yale University School of Medicine, New Haven, Connecticut, USA; Yale Center for Molecular and Systems Metabolism, Yale University School of Medicine, New Haven, Connecticut, USA; Current address: Boston Consulting Group, 466 Springfield Ave, Summit, NJ, 07901

**Keywords:** Mitogen-activated protein kinase (MAPK), MAPK phosphatase (MKP), allostery, NMR spectroscopy, molecular dynamics and X-ray crystallography

## Abstract

Dual-specificity mitogen-activated protein kinase (MAPK) phosphatases (MKPs) directly dephosphorylate and inactivate the MAPKs. Although the catalytic mechanism of dephosphorylation of the MAPKs by the MKPs is established, a complete molecular picture of the regulatory interplay between the MAPKs and MKPs still remains to be fully explored. Here, we sought to define the molecular mechanism of MKP5 regulation through an allosteric site within its catalytic domain. We demonstrate using crystallographic and NMR spectroscopy approaches that residue Y435 is required to maintain the structural integrity of the allosteric pocket. Along with molecular dynamics simulations, these data provide insight into how changes in the allosteric pocket propagate conformational flexibility in the surrounding loops to reorganize catalytically crucial residues in the active site. Furthermore, Y435 contributes to the interaction with p38 MAPK and JNK, thereby promoting dephosphorylation. Collectively, these results highlight the role of Y435 in the allosteric site as a novel mode of MKP5 regulation by p38 MAPK and JNK.

## Introduction

The mitogen-activated protein kinases (MAPKs) are a family of serine-threonine kinases that are responsible for a variety of critical cellular functions^1^. The MAPKs are regulated in a well-controlled manner through phosphorylation by upstream kinases known as MAPK kinases (MKKs), which activate the MAPKs^2, 3^. In contrast, the MAPKs are negatively regulated via dephosphorylation by enzymes known as MAPK phosphatases (MKPs)^4, 5^. The coordinated activities between the MKKs and MKPs dictate the signaling output of the MAPKs to propagate responses such as gene transcription^4, 6^. Although there has been a substantial amount of work conducted on the actions of the MKKs, such as MEK1/2^7^ on MAPK activation, much less is known about the regulation of the MKPs on MAPK inactivation. Given the increasing realization that the MKPs represent novel therapeutic targets for the treatment of cancer, inflammatory and metabolic diseases, as well as rare diseases^8, 9, 10^, a detailed understanding of MKP regulation has gained in its importance and significance.

The MKPs are a group of ten dual-specificity phosphatases (DUSPs) that directly inactivate the MAPKs through the dephosphorylation of the regulatory phosphotyrosine and phosphothreonine residues that reside within the activation loop of the MAPKs^11^. The MKPs exhibit an exceptionally high degree of specificity for the MAPKs via direct interactions with the MAPKs through a conserved N-terminal MAPK docking site designated as the kinase interacting motif (KIM)^12^. Further selectivity to the MAPKs is conferred through a well-conserved catalytic pocket at the C-terminus comprising the essential cysteine residue. The structure of the catalytic pocket is sufficiently deep to accommodate the phosphotyrosyl residue of the MAPK to coordinate the first dephosphorylation event, followed by phosphothreonyl dephosphorylation^13, 14^. In addition to the combined contributions of the KIM and the active site in establishing MKP-MAPK dephosphorylation specificity, the MKPs typically reside in a low basal state of activity and often adopt an active conformation upon binding MAPK^14, 15, 16^. Thus, the MKPs are highly regulated and direct exquisite MAPK substrate specificity through multiple mechanisms.

Despite these well-defined modes of MKP regulation, we identified an additional regulatory mechanism of the MKPs that indicates that phosphatase activity can be regulated through a novel allosteric site within the catalytic domain^17^. A high-throughput screen identified Compound 1 (Cmpd 1) as a small molecule inhibitor of MKP5 catalytic activity, and mutational analyses in combination with a solved X-ray structure of the MKP5 catalytic domain (MKP5-CD) bound to Cmpd 1 revealed that Cmpd 1 interacts with MKP5-CD at an allosteric pocket approximately 8 Å away from the catalytic Cys408^17^. The interaction of the inhibitor with MKP5- CD was mediated through hydrophobic interactions, most notably via Tyr435, and disruption of allosteric pocket residues attenuated Cmpd 1-MKP5-CD binding and MKP5 catalysis^17^. The discovery of the MKP5 allosteric site thereby uncovered a novel mode of regulation for MKP5 catalysis. Indeed, the importance of the MKP5 allosteric site for catalysis is supported by the observation that the critical Tyr435 residue is conserved amongst the active MKPs. We showed that MKP7, which is the nearest MKP in sequence homology to MKP5, contained an analogous allosteric site within its catalytic domain^18^. The MKP7 allosteric site is not only critical for MKP7 catalysis, but also contributes to binding with its substrates, p38 MAPK and c-Jun N-terminal kinase (JNK)^18^. These observations expanded the possible functions of the allosteric site, suggesting a role in not only regulating MKP catalysis but also MAPK engagement.

To further define the molecular characteristics and function of the MKP5 allosteric site, here we performed a combination of biophysical and biochemical experiments. We report that the MKP5 allosteric site confers profound conformational changes consistent with our previously reported deleterious effects of mutations in this region on MKP5 catalysis^17^. Molecular dynamics simulations infer that the allosteric site represents a conduit through which conformational dynamics are mediated, and together with NMR and X-ray crystallography, provide a plausible explanation for the impact of the allosteric site on MKP5 catalytic activity. Finally, we show that the MKP5 allosteric site represents an essential region through which the stress-activated MAPKs, p38 MAPK and JNK, interact to mediate dephosphorylation. Together, these data shed new light on the molecular and functional characteristics of the MKP5 allosteric site and provide further information on how selectively targeting this allosteric site within MKP5, and possibly other MKPs, leads to enzymatic inhibition.

## Results

### Inhibitor-bound MKP5-CD crystal structures exhibit conformational changes in both the allosteric and catalytic pockets

In our previous study using high throughput screening, we identified Cmpd 1 as an inhibitor of MKP5 catalytic activity. We determined the structure of the complex between MKP5-CD and Cmpd 1, characterized the kinetics of Cmpd 1-mediated inhibition of MKP5 activity which led to its assignment as an allosteric MKP5 inhibitor. The key allosteric residue that mediates Cmpd 1 binding, Y435, was identified to be essential for MKP5 catalysis^17^. To further define the molecular basis through which Y435 regulates MKP5 function we reasoned that structures of mutations of this allosteric residue in the presence and absence of Cmpd 1 would reveal further insights into the subtleties of the allosteric pocket. Since the variant Y435W retains 70% of WT MKP5-CD activity and binds Cmpd 1, whereas the Y435A and Y435S variants were catalytically impaired and failed to bind Cmpd 1, we focused on solving the structure of Y435W in the presence and absence of Cmpd 1. Crystals of the apo Y435W-CD structure and the Cmpd 1-bound complex were solved at 2.40 and 2.90 Å, respectively (Supplementary Table 1). These two new structures were compared to (a) each other, (b) to apo-WT MKP5-CD (PDB: 2OUD)^19, 20^, and (c) to Cmpd 1-WT MKP5- CD (PDB:6MC1)^17^ for the overall total RMSD (Fig. 1). Structural alignment of the allosteric residue in the apo-form indicates the plane of the tryptophan in Y435W has tilted 42.8° away from the tyrosine in WT MKP5-CD (Fig. 1a). The electron density of Cmpd 1 is fully interpretable in Y435W (Supplementary Fig. 1). In the Y435W-Cmpd 1 complex, W435 is tilted 64.9° and shifted toward the α4 helix, creating space for Cmpd 1. No major differences were observed between the Cmpd 1-bound WT and Y435W complex structures (Fig. 1a). Additionally, multiple hydrophobic interactions are established between A416, I420, I445, N448, and P451 and the dimethyl butanone moiety of Cmpd 1. The backbone atoms of M452 and N448 also serve as electron donors for hydrogen bonding with Cmpd 1 (Fig. 1b).

**Figure 1.**
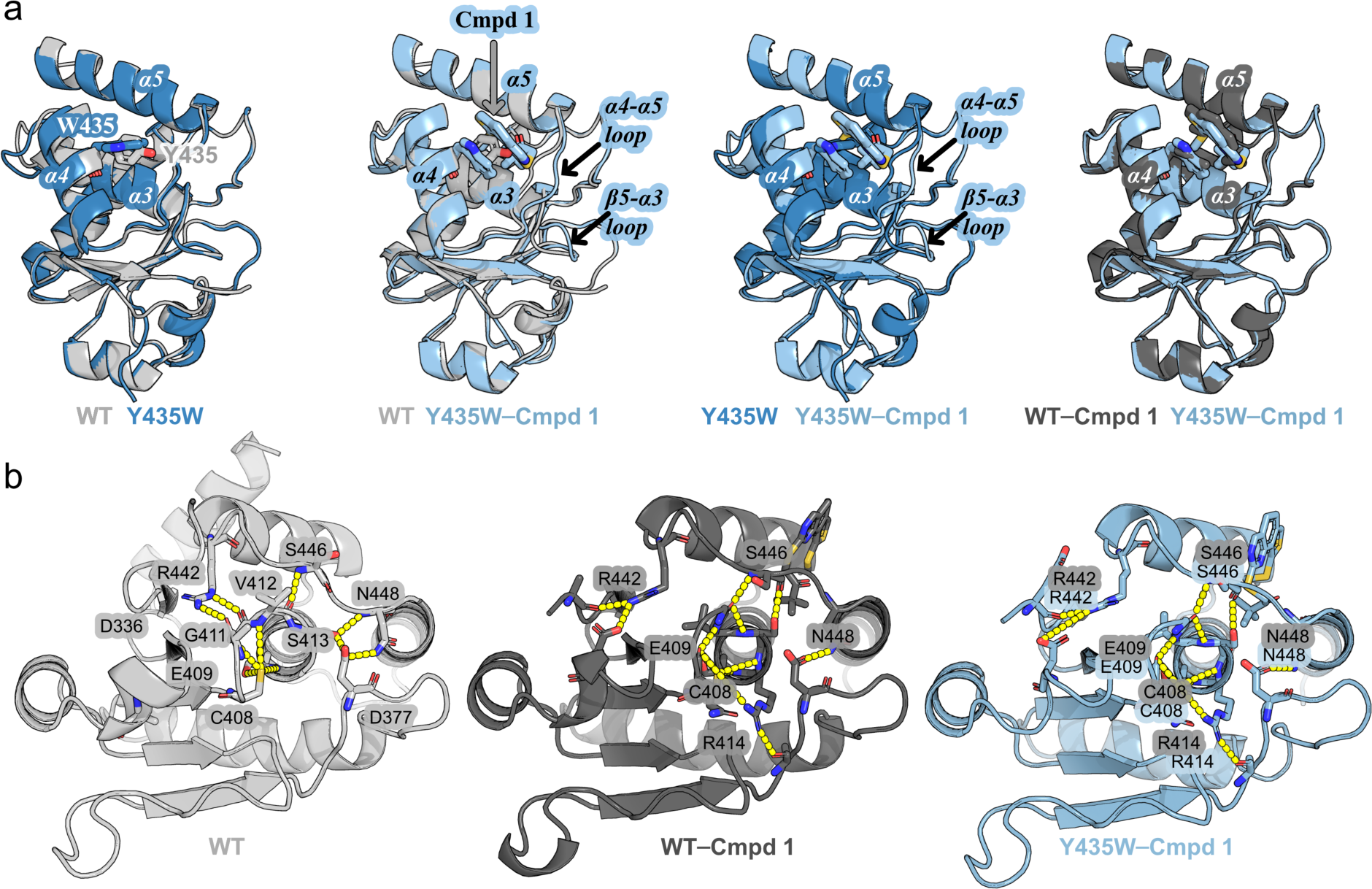
Tryptophan mutation at residue 435 confers structural and dynamic changes to the MKP5 allosteric pocket. **(a) Left to right,** structural overlay of the catalytic domain of WT MKP5 (light gray) and Y435W (marine blue), WT MKP5 (light gray) and Cmpd 1-bound Y435W (light blue), Y435W (marine blue) and Cmpd 1-bound Y435W (light blue) and WT-Cmpd-1 bound (dark gray) overlayed with Y435W-Cmpd-1 bound (light blue). The structural differences in the *α4-α5* loop and the catalytic pocket are labeled. **(b)** Network of hydrogen bonds that connects loop *α4-α5* loop of the allosteric pocket to the catalytic pocket in the apo WT MKP5 (**left**), and the dynamic changes in the secondary structure and new hydrogen bonds formed upon Cmpd 1 binding to WT MKP5 **(middle)** and to Y435W MKP5 **(right).**

The Cmpd 1-WT MKP5-CD structure showcases alterations in the conformation of *α4-α5* loop and the catalytic pocket that reduces the volume of the enzymatic site by ∼20%^17^. There are some notable differences to account for the reduction, particularly the movement of loop residues (I445-N448) in the presence of Cmpd 1 that alters the hydrophilic interactions between the residues of *α4-α5* loop (allosteric pocket) and *α3* helix (catalytic pocket) (Fig. 1b). This observed change includes the side chain of S446 and P447 on the loop between *α4-α5*, and a slight rotation of helix α3, resulting in rearrangements distinct from apo MKP5-CD. Other structural modifications involving helix *α3* are particularly important because its N-terminal region (residues G411-A416) separates the allosteric and enzymatic sites. A new hydrogen bond is formed between the backbone carbonyl of S446 and the hydroxyl group of S413 in the *α3* helix by the rearrangement caused by Cmpd 1. These new interactions probably occur concurrently with the disruption of a hydrogen bond between the backbone atoms of S413 and N448. A new hydrogen bond between the backbone amide of S446 is made with the carboxyl of G411 after the disruption of the bond between S446 with V412. In the apo form, R442 interacts and stabilizes A410 and G411, but these interactions disappear and promote the conformational changes of the *β5-α3 loop* in the catalytic pocket. Significantly, this promotes a new interaction between catalytic residues C408 and Q409. Finally, the tilt between the Y435 and W435 in the Cmpd 1-bound structures contributes to changes in the allosteric and catalytic pockets. We explored whether the ∼70% enzymatic activity of Y435W (relative to WT MKP5) provided a sufficient difference in the volume of the catalytic site, but it was not significant. Although the two other mutations, Y435A and Y435S, which have little enzymatic activity and do not bind Cmpd 1^17^, are well-suited for studying the structural and mechanistic source of catalysis, they were unstable for crystallization. Hence, we studied the effect of the Y435A and Y435S mutations on the apo proteins by NMR and molecular dynamics (MD) and focused on WT and Y435W for the Cmpd 1 bound proteins.

### Aromaticity of residue 435 is required for structural fidelity of the allosteric pocket

NMR studies were performed to characterize the chemical importance of the allosteric pocket residue Y435 by determining the structural effects of mutations at that allosteric site. Variants at Y435W, Y435S, and Y435A were generated, and all were found to be suitable for these NMR studies. Analysis of ^1^H^15^N HSQC correlation spectra revealed chemical shift perturbations indicating localized structural changes, and line broadening suggesting heightened dynamics, at residues surrounding the site of mutation (Fig. 2a, b and Supplementary Fig. 2). Though such structural and dynamic effects are often observed in regions adjacent to a mutation, the Y435S and Y435A variants exhibit considerably more chemical shift and dynamic perturbations than Y435W. These data suggest that the loss of side-chain aromaticity at residue 435 disrupts the integrity of the allosteric pocket, which is consistent with previous work showing reduced binding of Cmpd 1 to MKP5 when Y435 is mutated to either a serine or alanine, while binding and activity are maintained by a tryptophan residue at the same site^17^. Further, resonances exhibiting chemical shift perturbations in Y435S and Y435A often shift on a similar trajectory relative to WT MKP5-CD. These observations suggest that the structural changes induced by Y435S and Y435A populate a similar conformation, which is distinct from WT MKP5-CD (Fig. 2c). In contrast, Y435W induces chemical shift perturbations at those same resonances, but on a divergent trajectory to that of the serine and alanine variants or remaining similar to WT MKP5-CD. Notably, all three Y435 mutations result in a chemical shift perturbation at the catalytic residue D377, demonstrating that disruption of the allosteric site through mutation is propagated to the catalytic site, as expected for an allosteric system. These data support previous findings that Y435 mutations decrease MKP5 activity, with the degree of NMR-detected perturbation directly correlating with the degree of catalytic attenuation^17^. Similarly, thermal unfolding experiments revealed that the Y435S and Y435A mutations cause significant destabilization of MKP5-CD, with changes in the melting temperature (Δ*T*_m_) of -11.9°C and -14°C, respectively, relative to WT (Supplementary Fig. 3). Conversely, Y435W only exhibits a Δ*T*_m_ of -4°C. The rank order of changes in the melting temperature also correlates with the variant catalysis^17^. Thus, in addition to the integrity of the allosteric pocket, the aromaticity of the side chain at residue 435 contributes to the global stability of the MKP5-CD and further implicates the allosteric pocket as critical for MKP5 structural integrity.

**Figure 2.**
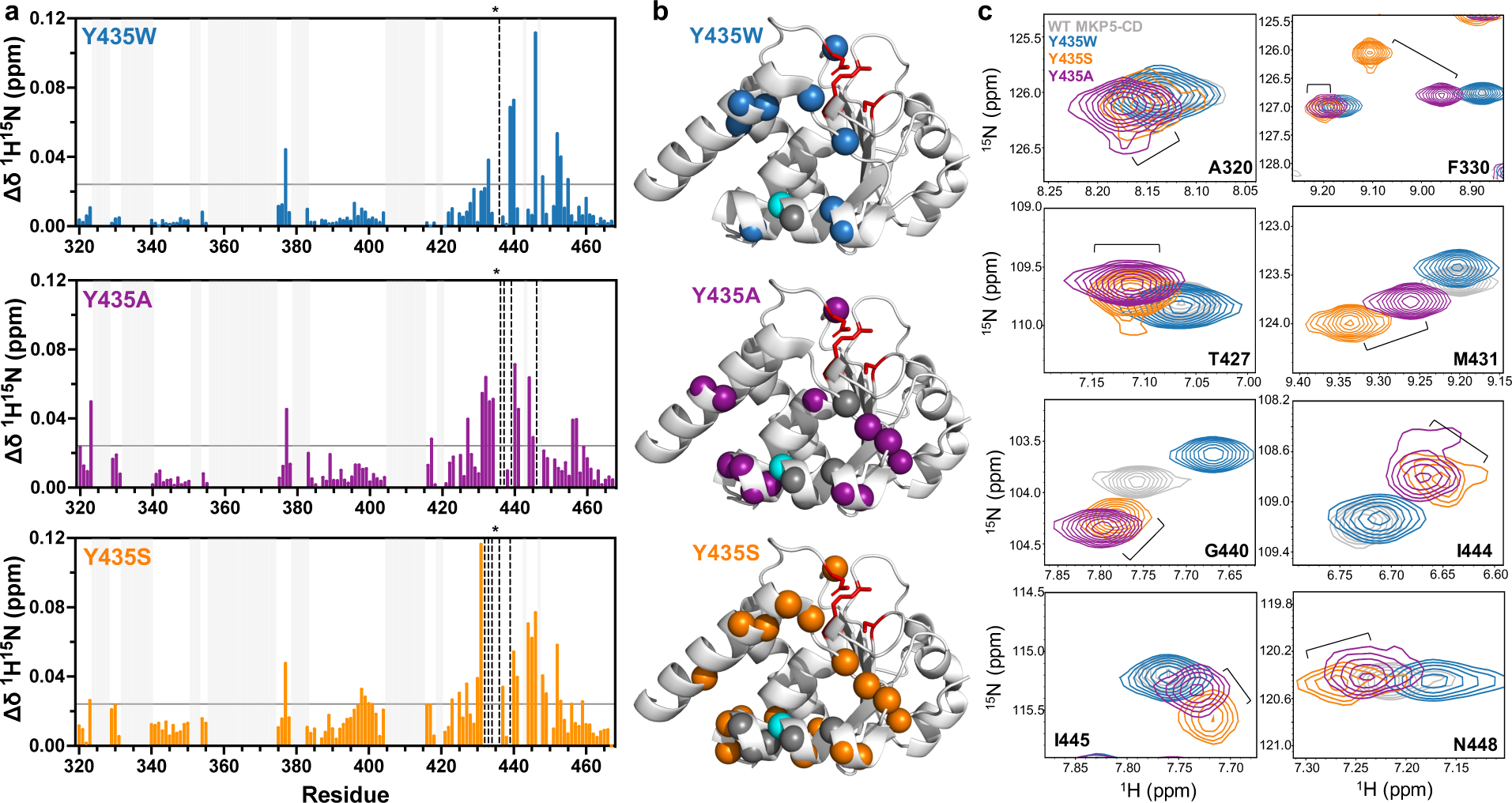
Loss of aromaticity at residue 435 elicits structural and dynamic perturbations in the allosteric pocket. **(a)** Chemical shift perturbation plots for Y435W (blue), Y435A (purple), and Y435S (orange) MKP5-CD. Colored bars represent ^1^H^15^N combined chemical shift perturbations (Δδ), calculated for each mutant relative to WT MKP5-CD. Significant chemical shift perturbations were determined as those 1.5σ above the 10% trimmed mean of all data sets (gray horizontal line). Black vertical dashed lines denote resonances that exhibit line broadening, and gray vertical lines indicate residues that are unassigned. The site of mutation is noted by an asterisk above each plot. **(b)** Significant chemical shift perturbations are plotted onto the MKP5-CD structure (PDB: 2OUD) in blue (Y435W), purple (Y435A), and orange (Y435S) spheres. Line-broadened resonances are indicated by gray spheres, the site of mutation is denoted by a cyan sphere, and the side-chains of residues at the catalytic site (D377, C408, R414) are highlighted in red. **(c)** Representative snapshots from the ^1^H^15^N HSQC spectral overlay of WT MKP5-CD (gray), Y435W (blue), Y435S (orange), and Y435A (purple) demonstrate that the Y435S and Y435A mutants exhibit similar chemical shift trajectories, suggesting a similar conformation populated by those mutants, while Y435W chemical shifts diverge from that of the serine and alanine mutants and often resemble WT.

### Molecular dynamics identifies MKP5 allosteric site-to-active site communication channel

To gain further insight into the impact of the allosteric site mutations, we analyzed differences in root mean squared fluctuations (ΔRMSF) from MD simulations. Significant mutation-induced changes in the vicinity of the MKP5 allosteric site and the flexible regions encompassing residues 325-350 and 353-379 were noted (Fig. 3a, b), which remained unassigned in the NMR chemical shift profiles. In general, the RMSF patterns by MD mirror the trends seen in the chemical shift and dynamic perturbations, with the greatest fluctuations detected in the serine variant (Fig. 3a, b).

**Figure 3.**
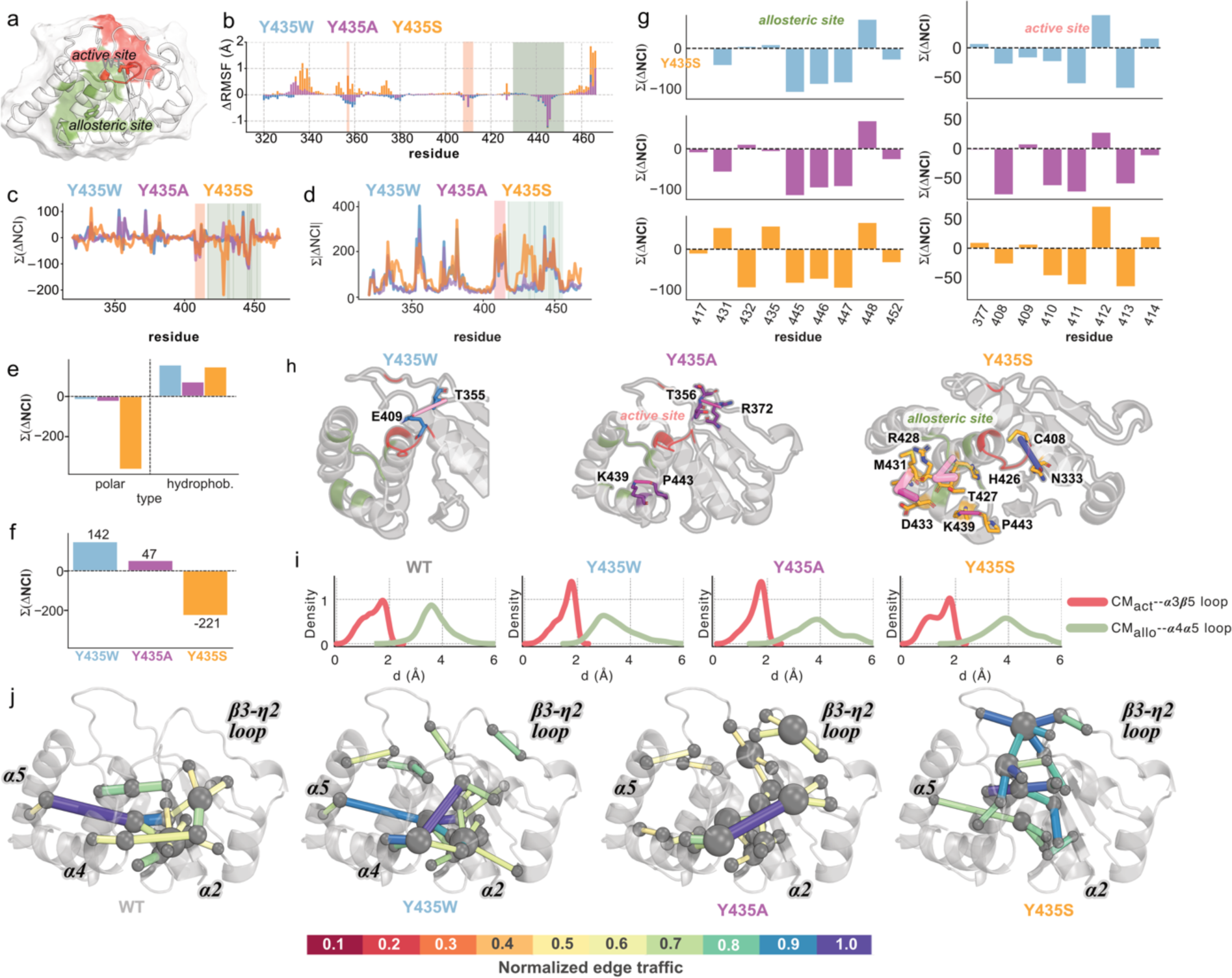
Loss of aromaticity at the allosteric residue 435 reorganizes non-covalent interactions. (**a**) Cartoon representation of MKP5 highlighting the active (red) and allosteric site (green). (**b**) Difference RMSF (ΔRMSF) of backbone fluctuations with respect to those sampled in WT MKP5, computed over 3 µs trajectories for Y435W (blue), Y435A (purple), and Y435S (orange). (**c, d**) Non-covalent interaction (Σ(ΔNCI) and Σ|ΔNCI|) difference profiles computed for each variant by taking the cumulative persistence of non-covalent interactions sampled for each residue (using 1 frame every 5 ns) and taking the difference with respect to the WT state. **(e, f**) Cumulative ΔNCI profiles for all residues, showing individual polar and hydrophobic contributions (e) or both (f). (**g**) Cumulative ΔNCI contributions for residues in the allosteric and active sites. All ΔNCI are sampled along 3 µs of molecular dynamics trajectories in each state, using one frame every ns. **(h)** Selected ΔNCI changes (relative to WT) with an overall persistency greater than 15% that are unique to each mutant, highlighting the regions that are most susceptible to the introduction of each mutation. **(i)** In red, distribution of distances computed between the active site center-of-mass (CM) and heavy atoms in the *β5-α3* loop. In green, distribution of distances computed between the allosteric site center-of-mass and heavy atoms in the *α4-α5* loop. Centers-of-mass is computed using heavy atoms in the active site (residues 408-414) and in the allosteric-site (residues 442-452). **(j)** Traffic maps show the preferential communication routes in each state.

We quantified changes in non-covalent interactions (ΔNCI) observed in MD simulations of the MKP5-CD variants relative to the WT MKP5-CD trajectory. The unperturbed interaction network of WT MKP5-CD is illustrated in Supplementary Fig. 4. We focused on the reorganization of interactions during microsecond-scale MD, which were performed in triplicate to ensure sufficient sampling and to enable comparison with experimental data. Figures 3c and 3d depict the cumulative changes in non-covalent interactions per residue, denoted as Σ(ΔNCI), and the absolute changes, denoted as Σ|ΔNCI|, arising from polar and apolar interactions. The Σ(ΔNCI) metric offers an assessment of the overall alteration to the chemical environment at each residue. While the absolute value, Σ|ΔNCI|, highlights the residues experiencing the most significant alterations in the NCI network, irrespective of whether the net change is zero (i.e. in cases where disrupted interactions are compensated by the formation of new ones). Significant deviations from the WT MKP5-CD profile are observed, regardless of the mutation at residue 435, with the most notable changes concentrated in the active and allosteric sites (Fig. 3c, d). The Y435S variant exhibits a distinct Σ|ΔNCI| profile, especially in the allosteric site and at residues 334-344 (Fig. 3d). The profile of Σ|ΔNCI| indicates similarly reshuffled interactions caused by the Y435W and Y435A mutations, although there are significant differences between their relative intensities (Fig. 3d-f). The net change induced by each variant follows the trend Y435W > Y435A > Y435S, suggesting that the increasing disruption of NMR chemical shifts caused by non-aromatic mutations corresponds to a greater loss of non-covalent interactions induced by the variants. This loss is accounted for largely by polar interactions in Y435S, while the increase of ΔNCI in Y435W is mostly derived from an increase in hydrophobic contacts (Fig. 3e). Such changes include the disruption of the catalytic residue C408:SG–G411:O hydrogen bond, which decreases by over 35% with Y435W, Y435A, and Y435S mutations. Conversely, the N448:N–L449:N backbone hydrogen bond shows a 30% increase with Y435W, Y435A, and Y435S mutations. The changes in ΔNCI values occurring in more than 15% of the analyzed trajectories for each variant are detailed in Supplementary Table 2. The Y435S mutation causes over 40% of the examined scenarios, including a rearrangement of the hydrogen-bonding network in the allosteric site (residues T427-N--R428-N, T430-OG1--D433-N, T430-OG1--M431-N), which is not observed in Y435W (Supplementary Table 2).

The cumulative ΔNCI contributions for residues in the allosteric and active sites suggest that the major perturbations occur in the allosteric site (Fig. 3g). However, the variants differ in the impact they have on the active site, as Y435S leads to a reinforcement of the hydrophobic contact C408-N333, which is not observed in either Y435W or Y435A (Table 1). Figure 3h shows selected interactions with an overall persistency greater than 15% that are unique to specific variants, highlighting the regions that are most susceptible to the introduction of each mutation. The largest difference between Y435W and Y435A is the disappearance of the T355:OG1- E409:NE2 hydrogen bond, unobserved in the latter, and concurrent loss of the K439-NZ--P433-O hydrogen bond in both in Y435A and Y435W, but not in Y435S (Fig. 3h). These results indicate that in comparison to WT MKP5-CD, Y435A and Y435S mutations enhance the flexibility of the active and alloseteric site, in contrast to the minor change observed for the tryptophan mutation, as shown in figure 3i. Here, we computed the distributions of the active site and allosteric site center-of-mass distances to *β5-α3* loop and *α4-α5* loop, respectively. These distances reflect the inner reorganization of these two functional sites, showing a shift of allosteric site distances to larger values in Y435A and Y435S (coupled to large changes in the active site in Y435S), which are not observed in WT and Y435W.

**Table 1.** The largest ΔNCI changes for each mutant (Y435W, Y435A, Y435S) with respect to the WT apo state. The selection includes the ΔNCI values relative to each mutant that differ from any other by more than 25%. All interaction values lower than 5% in absolute value are indicated as -. In the absence of measured ΔNCI, cells are left blank. Green and red colorings represent the allosteric and active site, respectively.

| Donor-Acceptor | $\Delta Y435W$ | $\Delta Y435A$ | $\Delta Y435S$ | Type | <u>Secondary Structure</u> |
| --- | --- | --- | --- | --- | --- |
| A435-OG--T432-N |  |  | 28.37 | polar | <u>$\alpha 4$</u> |
| **R372-NH1--T355-OG1** | 38.95 | 34.58 | - | polar | <u>$\beta 4$ -- $\beta 4$-$\eta 2$ loop</u> |
| R428-N--H426-N | - | - | -32.99 | polar | <u>$\beta 1$ -- $\beta 1$- $\beta 2$ loop</u> |
| R428-N--L423-O | - | - | -26.71 | polar | <u>$\beta 1$ -- $\beta 1$- $\beta 2$ loop</u> |
| R428-N--T427-N | - | - | -28.31 | polar | <u>$\beta 1$ -- $\beta 1$- $\beta 2$ loop</u> |
| **R442-NH1--L331-O** | 31.04 | 26.75 | - | polar | <u>$\alpha 4$- $\alpha 5$ loop -- - $\beta 2$</u> |
| **M431-N—T432-N** | - | -6.4 | 30.43 | polar | <u>$\alpha 4$</u> |
| T427-N—R428-N | - | 5.29 | -46.48 | polar | <u>$\alpha 3$-$\alpha 4$ loop</u> |
| **T430-OG1—D433-N** | -5.06 | -7.11 | 41.69 | polar | <u>$\alpha 3$-$\alpha 4$ loop -- $\alpha 4$</u> |
| **T430-OG1—M431-N** | - | - | -49.32 | polar | <u>$\alpha 3$-$\alpha 4$ loop</u> |
| **C408—N333** | - | - | 30.9 | hydrophobic | <u>$\beta 5$-$\alpha 3$ loop -- $\beta 2$ $\eta 1$ loop</u> |

To further understand how changes in the allosteric pocket affect conformational flexibility in surrounding loops and reorganize crucial catalytic residues in the active site, we computed a traffic map of residue-to-residue shortest and highest-correlation paths from the sampled dynamics of each mutant (Figure 3j). This traffic map includes paths for each residue pair, resulting in N² routes (where N is the number of residues), allowing us to estimate the "backroads" and "highways" of information transfer through the polypeptide (see Methods section for further details). We find similar traffic profiles in WT and Y435W, as opposed to remarkably altered communication pathways in Y435A and Y435S, each of which differs from WT in unique aspects (Fig. 3i). WT and Y435W show a high-traffic route connecting the allosteric region (via *α5*) with the active site through *α2*, suggesting that this is the preferred route of communication between allosteric and active sites (Fig. 3j). In contrast, the same connection is poorly sampled in Y435A, which points to an inefficient coupling between the two functional sites (Fig. 3j). Y435S completely rewires the communication routes, consistent with the experimentally observed trends of strong NMR chemical shift perturbations (Fig. 2b) and poor catalytic activity^17^. Residues that exhibit structural perturbations in Y435A (L456 and E457) and Y435S (E459) are in close proximity to residues on *α5* which is implicated in the preferential allosteric-to-active site communication routes for WT Y435W (F458 and D461). These data suggest that the structural changes induced by the Y435A and Y435S mutations at these *α5* sites contribute to the altered dynamics of neighboring residues, causing an attenuation of communication via the *α5*-*α2* route to ultimately impact catalytic behavior.

### Allosteric site ligand engagement changes dynamics and non-covalent interactions in the MKP5 catalytic domain

Biochemical studies of MKP5 in the presence of Cmpd 1 revealed that mutation of Y435 to serine or alanine dramatically reduced its ability to bind^17^. In contrast, the Y435W variant showed slightly reduced, but near-WT affinity^17^. We investigated the interaction of Cmpd 1 with MKP5-CD by NMR and found that when Cmpd 1 is bound to either WT MKP5-CD or Y435W, the associated ^1^H^15^N HSQC NMR spectra exhibit significant line broadening (Fig. 4a, b and Supplementary Figs. 5 and 6). This dynamic effect, seen across the entire domain, implies that Cmpd 1 when bound to the allosteric site exerts its inhibitory effect by disrupting active-to- allosteric site communication via micro-millisecond dynamics. Such extensive line broadening was surprising for small molecule binding and could also occur due to the formation of large oligomers. However, size exclusion chromatography suggests that the signal attenuation observed by NMR is not due to a change in the oligomeric state of MKP5-CD (Supplementary Fig. 7). Conversely, titration of Cmpd 1 into Y435S produced minor NMR spectral changes (Supplementary Figs. 8), consistent with negligible binding and prior biochemical findings that can be attributed to structural and dynamic disruption of the allosteric pocket (Figs. 2 and 3) and loss of π-stacking interactions between Cmpd 1 with an aromatic side chain at residue 435^17^.

**Figure 4.**
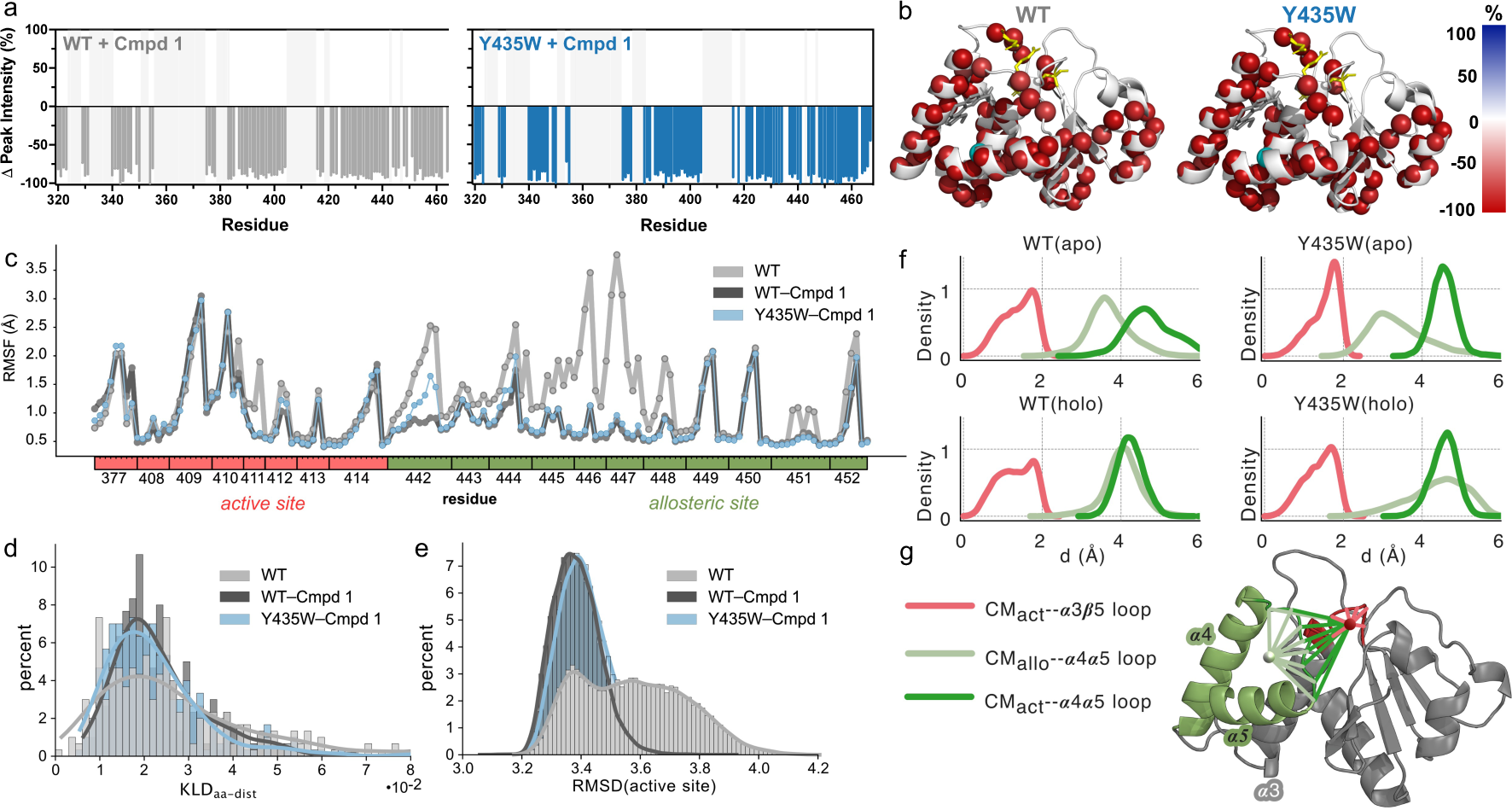
Allosteric site engagement alters the dynamic profile of the MKP5-CD. **(a)** Changes in ^1^H^15^N HSQC resonance (peak) intensity when MKP5 is bound to Cmpd 1 at the allosteric site, calculated relative to apo MKP5 for WT (gray) and Y435W (blue). Light gray vertical lines on all plots indicate residues that are unassigned. **(b)** Δ peak intensity is plotted onto the MKP5 structure (PDB: 6MC1), with blue and red spheres representing decreases and increases in resonance intensity, respectively, and the color gradient denoting the magnitude of the change in resonance intensity in the presence of Cmpd 1 relative to apo MKP5. Residue 435 is represented by a cyan sphere, and the catalytic side-chains are highlighted in yellow. Cmpd 1 is modeled into the allosteric pocket for reference. **(c)** Dynamics of WT, WT-Cmpd 1, and Y435W-Cmpd 1 shown through Kernel density estimates of the Kullback-Leibler (KLD) divergence distributions of WT (apo; gray), WT (holo; dark-gray), and Y435W (holo; light-blue) MD-sampled configurations with respect to the crystal structure (6MC1). The KLD was computed using the heavy atoms belonging to the residues in the panel, namely the active site and *α4-α5* loop. **(d)** Kernel density estimates of RMSD distributions of WT (apo), WT (holo), and Y435W (holo), computed on the active site and allosteric site heavy atoms. **(e)** RMSF distributions of WT (apo), WT (holo), and Y435W (holo), computed on active site and α4-α5 loop. **(f)** In red, distribution of distances computed between the active site center-of-mass and heavy atoms in the *β5-α3* loop. In light green, distribution of distances computed between the allosteric site center-of-mass and heavy atoms in the *α4-α5* loop. In dark green, distribution of distances computed between the active site center-of-mass and heavy atoms in the *α4-α5* loop. Centers-of-mass are computed using heavy atoms in the active site (residues 408-414) and in the allosteric-site (residues 442-452). **(g)** MKP5-CD structure showing the allosteric site (green), active site (red), and the centers-of-mass of each site (light-green and red spheres, respectively). Allosteric site center-of-mass to *α4-α5 loop* Cα-atom distances are shown in light-green lines. Active site center-of-mass to *α4-α5 loop* Cα-atom distances are shown in dark-green lines. Active site center-of-mass to *β5-α3 loop* Cα-atom distances are shown in red lines.

Based on these experimental findings, we simulated the Cmpd 1-bound dynamics for WT and Y435W MKP5-CD. The stability of Cmpd 1 within the Y435W binding site resembles that of the WT. In both complexes, Cmpd 1 adopts a specific configuration supported by a strong π- stacking interaction between the aromatic portion of the ligand and tyrosine or tryptophan residues (Supplementary Fig. 9). The modified aromatic ring portion leads to a slightly increased flexibility of the Y435W binding site (Supplementary Fig. 10).

A comparison of the MD-derived conformations of the *α4-α5* loop of the allosteric and active sites showed remarkable alignment (Figs. 4c-e). To characterize the difference in the conformational dynamics among apo and WT or Y435W Cmpd 1-bound structures, we measured the similarity of the MD configurations and crystal structure (6MC1) using the Kullback-Leibler divergence (KLD) metric computed at each frame of the trajectory. The probability distribution computed from the resulting values shows higher similarity between the WT and Y435W holo distributions (narrower Gaussian centered on smaller KLD values) when compared to the apo WT (Fig. 4d, e). These results agree with those derived from the crystal structure wherein a comparison of the holo WT and holo Y435W indicates little difference between the structures (Fig. 1). Moreover, the distance distributions between active site, allosteric site, and relevant loops (*a4-a5* loop*, β5-α3* loop) are conserved in Y435W with the allosteric site center-of-mass (COM) to *a4-a5* loop distances showing a shift to larger distances, and active site COM to *β5-α3* loop distances distribution becoming narrower in response to Cmpd 1 binding (Fig. 4f-g), consistent with an active site volume reduction^17^.

We evaluated the impact of Cmpd 1 on WT and Y435W MKP5-CD by assessing the cumulative response based on NCI alterations that occur upon binding. These changes represent a range of persistency levels sampled at various frequencies throughout the trajectory. In both systems, the overall change in NCIs (ΔNCI (holo – apo)) in WT and Y435W results in an apparent "loss" of interactions (Supplementary Figs. 11, 12). This suggests a higher number of non-covalent interactions are being strengthened rather than weakened, indicating a propensity for continuous partner exchanges. This behavior is illustrated by the specific changes in ΔNCI (holo – apo), such as polar, hydrophobic, and π-stacking interactions on the WT protein structure (Supplementary Fig. 11). In the WT MKP5-CD, Cmpd 1 binding induces widespread modifications across the entire structure, particularly affecting the allosteric site (*α4*, *α5*), the active site (*β5-α3* loop), and neighboring secondary structures (*β5-η2* loop, *α1*, *η2*, *β4* – Supplementary Fig. 12). A similar pattern is observed in Y435W, albeit with a noticeable decrease in the secondary structures at *β3* and *β5* accompanied by more pronounced changes in *β5-η2* loop, *α1*, *η2*, and *β4* (Supplementary Fig. 12). The reorganization of these interaction networks (with respect to the apo states) is consistent with the significant line broadening observed in NMR experiments (Fig. 4a, b) and support the notion that Y435 is required to maintain the structural integrity of the allosteric pocket. The dynamic effects introduced by Cmpd 1 are consistent with the altered catalytic output, though the differences in overall active-allosteric site atomic displacements and non-covalent interactions in WT and Y435W mirrors the Cmpd 1-induced functional effect in the two systems.

### MKP5 allosteric site residue Y435 mediates MAPK binding

Previously, we showed that Y435 serves as a critical determinant of the allosteric pocket that regulates MKP5 catalysis^17^. Furthermore, structural insights from our NMR, crystallographic and MD studies demonstrated that disruption of the allosteric site either by mutation or Cmpd 1 binding caused disruption of loops, which appeared important for the transmission of structural changes to the active site. Furthermore, molecular modeling of Cmpd 1 bound to MKP5-CD with JNK1 predicts that an interaction interface is disrupted by Cmpd 1. These observations suggest that the allosteric pocket could serve to mediate MAPK interactions with MKP5-CD leading to changes in active site conformation. Based on these observations, we tested the hypothesis that the functional importance of these structural changes is evoked by the engagement of MKP5 with its substrates, p38 MAPK and JNK. To test this, we transiently transfected COS-7 cells with GST-tagged full-length WT MKP5 or MKP5 allosteric pocket variants, Y435W, Y435A and Y435S. As a control, COS-7 cells were also transfected with the catalytically inactive and substrate-trapping variant of MKP5 containing an active site Cys408-to-Ser mutation (C408S). To test for p38 MAPK or JNK binding to these MKP5 variants, either HA-tagged p38 MAPK or GFP-JNK expression vectors were co-transfected, respectively. Following affinity precipitation of MKP5 from COS-7 cells, we found that Y435A, Y435S, and Y435W MKP5 were significantly impaired in their ability to complex with p38 MAPK as compared with WT MKP5 (Fig. 5a). Similarly, JNK was also reduced in its amount that was complexed with the MKP5 allosteric pocket variants, as compared with WT MKP5 (Fig. 5b). In contrast, the catalytically inactive and substrate trapping variant C408S bound to both p38 MAPK and JNK at substantially higher levels, consistent with the establishment of a dead-end MKP5-JNK and MKP5-p38 MAPK complex. These results indicate that the MKP5 allosteric pocket not only mediates catalysis, as demonstrated previously^17^, but also mediates both p38 MAPK and JNK binding. The implications of MAPK docking to the allosteric site based collectively on our structural findings described herein infer that this interaction likely induces similar conformational changes that propagate to the structural reorganization of the active site.

**Figure 5.**
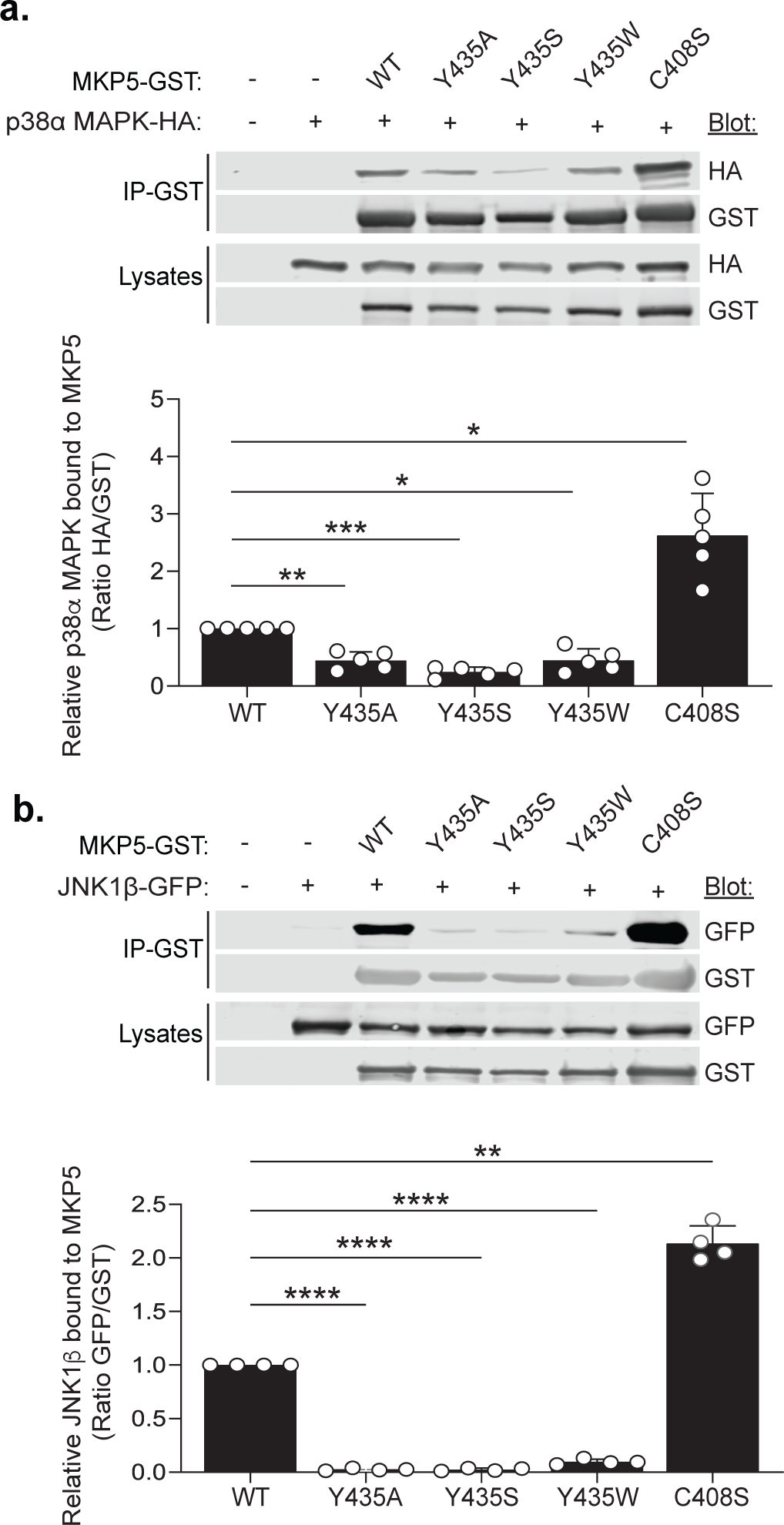
MKP5 residue Y435 mediates MAPK binding. Full-length GST-tagged MKP5 WT, Y435 mutants, and C2408S were transiently transfected with p38α MAPK-HA, JNK1-β-GFP, or ERK2-HA into COS-7 cells for 48 h. Cells were harvested and lysed, followed by affinity purification of GSH Sepharose beads. **(a)** GST affinity complexes and whole cell lysates were immunoblotted with anti-HA and anti-GST antibodies. **(b)** GST affinity complexes and whole cell lysates were immunoblotted with anti-GFP and anti-GST antibodies. Graphs shown below represent quantitation of **a**, p38α MAPK-HA/GST-MKP5 and **b**, JNK1-GFP/GST-MKP5. Data represent the mean ± SEM of three to four to five independent experiments. Statistical significance shown was generated using a two-way ANOVA. Key: *; *p*<0.05, **; *p*<0.01, ***; *p*<0.005 and *p*<0.0001.

### MKP5 allosteric site differentially orders p38 MAPK and JNK active site engagement

To further understand the relationship between MAPK-MKP5 allosteric and active site engagement, we tested the effects of the catalytically inactive substrate-trapping variant, C408S, to bind either p38 MAPK or JNK in the context of the allosteric Y435S variant. As shown earlier when Y435S was expressed in COS-7 cells, both p38 MAPK and JNK failed to interact (Fig. 5). Consistent with the substrate-trapping properties of C408S, both p38 MAPK and JNK bound with significantly increased levels to this MKP5 variant as compared with WT MKP5 (Fig. 6). However, when both the allosteric site (Y435A) and catalytically inactive substrate-trapping (C408S) variants were combined to generate the double Y435A/C408S MKP5 mutant, we found that p38 MAPK was still capable of complex formation with MKP5 (Fig. 6a-c). In contrast, JNK binding to MKP5 was markedly inhibited in the Y435A/C408S variant (Fig. 6d-f). Based upon our NMR findings that mutation of Y435 to A435 causes structural and dynamic perturbations in *α4* and *α5* regions of the allosteric pocket, these results suggest that JNK utilizes these regions to bind MKP5 more so than p38 MAPK. The levels of phosphorylated p38 MAPK and JNK present in the complex of Y435A/C408S MKP5 were reduced slightly for p38 MAPK and JNK as compared with C408S MKP5 alone (Fig. 6). We interpret these results to suggest that the active site conformational changes induced by Y435A at C408 are more deleterious to the formation of the phospho-thioate intermediate of phosphorylated p38 MAPK as compared with phosphorylated JNK. Collectively, these data suggest that p38 MAPK and JNK exhibit distinct differences in the constraints required for active site engagement when bound to the allosteric pocket.

**Figure 6.**
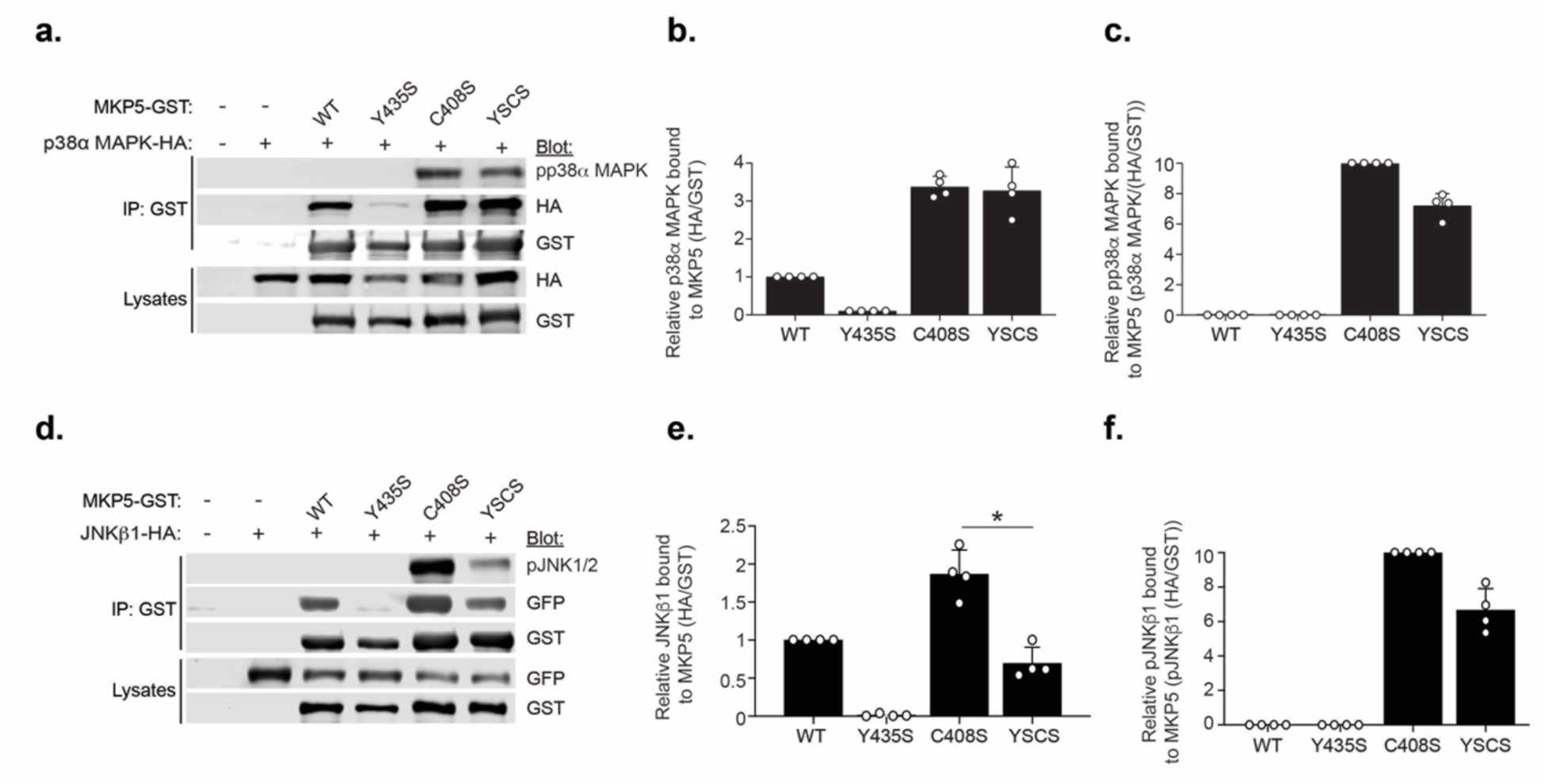
Effects of MKP5 residue Y435 on MAPK engagement at the active site. Full-length GST-tagged MKP5 WT, Y435A, C408S and C435/C408S mutants were transiently transfected with p38α MAPK-HA, JNK1-β-GFP into COS-7 cells for 48 h. Cells were harvested and lysed, followed by affinity purification with GSH Sepharose beads. **(a)** GST affinity complexes and whole cell lysates were immunoblotted with anti-HA, anti-GST and phospho-p38 MAPK antibodies. **(b)** Densitometric quantitation of relative normalized p38α MAPK bound to GST-MKP5; **(c)** Relative normalized phosphorylated p38 MAPK in complex with MKP5; **(d)** GFP affinity complexes and whole cell lysates were immunoblotted with anti-GFP, anti-GST and phospho-JNK antibodies. **(e)** Graph showing densitometric quantitation of relative normalized JNK bound to GST-MKP5; **(f)** Graph showing relative normalized phosphorylated JNK in complex with GST-MKP5. Data represent the mean ± SEM of four independent experiments. Statistical significance shown was generated using a two-way ANOVA. Key, *; p<0.05

## Discussion

The MKPs play fundamental and non-redundant signaling roles through their specific ability to dephosphorylate and inactivate the MAPKs in a spatio-temporal manner^4, 5^. Serving as nodal regulators of the MAPKs, the MKPs integrate multiple MAPK responses from upstream stimuli. As such, MKPs have been shown to mediate both positive and negative signaling outcomes and have become attractive therapeutic targets for certain human diseases^8^. Although the molecular basis of MKP’s ability to directly dephosphorylate the MAPKs is appreciated, how the MKPs derive signaling selectivity is not fully understood, suggesting that the full complexity of their regulation has yet to be uncovered. The identification of a novel allosteric pocket that regulates MKP5 activity uncovered an unanticipated mode of regulation that raised questions about the site-specific conformational changes at the molecular level that influence MKP5 catalysis. Here, we have integrated structural, computational, and biological approaches to demonstrate that the integrity of the allosteric site is strictly maintained through side-chain aromaticity at residue 435 located within the phosphatase domain. This interpretation is based on the conformation populated by the catalytically competent Y435W variant, visualized by crystallography and NMR, which is distinct from the dynamically unstable conformation of Y435S and Y435A. The Y435S and Y435A variants do not crystallize in the presence or absence of Cmpd 1. There are no major conformational changes between WT and Y435W structures in the presence or absence of Cmpd 1. The only minor change is the tilt of the bound Cmpd 1 in WT and Y435W, but this modification does not alter the volume of the active site, which is purported to be partly responsible for the decrease in enzymatic activity^17^. NMR and MD simulations highlight fluctuations near the allosteric site that are similar among the three variants, but also reveal that the distinct catalytic behaviors stem from a network of disrupted non-covalent interactions that follows a rank order of potency Y435W > Y435A > Y435S. Molecular conduits of communication between the allosteric and active sites interpreted by MD simulations provide insight into how ligand engagement propagates conformational changes to the active site. Finally, we show that both p38 MAPK and JNK represent physiological ligands of the allosteric site, suggesting that MKP5 undergoes dynamic conformational reorganization that likely facilitates MAPK dephosphorylation.

Although there have been other protein tyrosine phosphatases (PTPs) that have been reported to be modulated by allosteric small molecule binding, such as SHP2^21^ and PTP1B^22^, MKP5 represents the first MKP targeted in this manner^23^. MKP3/DUSP6 has been shown to be inhibited by the small molecule 3-(Cyclohexylamino)-2,3-dihydro-2-(phenylmethylene)-1*H*- inden-1-one (BCI) through a mechanism suggestive of allostery based upon *in silico* docking^24, 25^. However, how BCI, through this putative allosteric site, transmits its inhibitory effect to the active site remains unknown. Here, we have expanded upon our initial observation that Cmpd 1 collapses the active site volume to inhibit catalysis, by defining the communication channels from the allosteric site to the active site. Mutations at the allosteric site that deform hydrophobicity exert significant destabilization of the MKP5 catalytic domain. The Y435W and Y435A mutations impact the *β3-η2* loop, resulting in a restructuring of hydrogen bonds more notably disturbed in Y435S (see Supplementary Table 2). Within *β4-η3* loop, the catalytic residue D377 displays significant positional variation across all allosteric mutants, consistent with NMR chemical shift perturbations, with the interaction networks showing more pronounced changes in Y435A and Y435S. Thus, specific interactions affected by each variant *in silico* align with the observed experimental trends, supporting the interpretation that aromatic residues prime the MKP5-CD to facilitate its allosteric crosstalk. Thus, binding of Cmpd 1 to its allosteric site rewires non-covalent interactions (especially hydrogen bonds) and increases flexibility surrounding the pocket in a mutation-specific fashion. The distinct interaction networks observed from MD simulations with Cmpd 1 bound to MKP5 variants are also consistent with experimental findings.

Our biophysical findings, together with MAPK interaction assays, show that the allosteric pocket of MKP5 represents a multifunctional region within the catalytic domain to control enzymatic activity. Previous studies demonstrated that MKP5 exhibited a structure that is consistent with evidence that it has constitutive phosphatase activity^20, 26^. In other MKPs that have been reported to exhibit inducible activity, the catalytic aspartic acid residue is held away from the other catalytic residues by a hydrogen bond with a nearby asparagine, which is absent in MKP5^19, 20, 27^. The experiments reported here, which indicate the establishment of non-covalent networks from the allosteric site to the active site suggest that additional regulatory features of MKP5 are operative upon MAPK binding that could influence substrate selectivity, engagement and/or enhancement of phosphatase activity. Although both p38 MAPK and JNK interact with MKP5, they exhibit distinct modes of binding ^28, 29, 30^. p38 MAPK binds to the KBD of MKP5 whereas JNK utilizes a F/L-x-F motif within the catalytic domain of MKP5 ^30^. We now show that p38 MAPK and JNK differentially engage the active site cysteine since JNK fails to bind in context of the allosteric site/substrate-trapping mutant whereas p38 MAPK binding is retained. These results further support the observations of the distinct binding mechanisms of p38 MAPK and JNK to MKP5^19, 20, 28^. Indeed, these results imply that JNK binding to MKP5 proceeds first through its interaction with the allosteric pocket followed by catalytic site engagement. In contrast, p38 MAPK is retained by the substrate-trapping mutant in context of the allosteric site mutant, suggesting that the catalytic site engagement precedes that of allosteric site binding. Given that both p38 MAPK and JNK are predicted to alter catalytic site conformation, the extent to which allosteric site engagement regulates MKP5 activity will need to be determined. Furthermore, how p38 MAPK and JNK binding to this allosteric site is ordered in context of the KBD and Y/F-x-F motifs also requires further investigation. Nevertheless, it is noteworthy that the allosteric site is conserved amongst the active MKPs, including MKP7 where we have shown that the analogous allosteric site also contributes to both p38 MAPK and JNK binding^18^. Interestingly, MKP7 does not exhibit the observed differential binding with p38 MAPK and JNK via its active and allosteric site as seen for MKP5^18^. Thus, this allosteric site is likely to be a common site of MAPK binding amongst the MKPs but with differing contributions for MAPK binding relative to active site engagement.

MKP5 has been implicated as a potential therapeutic target for the treatment of fibrosis in skeletal muscle, heart and the lung ^31, 32, 33^. Given that there are limited treatments available for fibrotic tissue disease, the importance of understanding the molecular mechanisms of MKP5 allosteric inhibition is of significance. Collectively, this study provides molecular insight into the dynamic structural changes initiated upon ligand engagement of the MKP5 allosteric site. NMR and MD analyses reveal that mutations at the allosteric Y435 residue all result in disruption of the catalytic site at C408 and the general acid at D377 to varying degrees, providing direct evidence for allostery. MD simulations illuminate channels of communication between Y435 and the active site by critical non-covalent interactions. Finally, our data indicate that an additional feature of the MKP5 allosteric site is to bind MAPK. Collectively, these data uncover previously unappreciated modes of MKP5 regulation which can be harnessed to inform allosteric inhibitor design.

## METHODS

### Protein expression and purification

The WT and single-point mutants of the MKP5-CD were cloned into the pET28a vector with a TEV cleavage site and subsequently transformed into BL21 Gold (DE3) cells. Variant constructs were generated using Pfu Turbo DNA polymerase (Agilent). Protein expression and purification followed our previously reported protocol^17^, with minor modifications. Bacterial cells were induced using 1 mM IPTG and allowed to express the protein overnight at 20 °C. The cells were suspended in lysis buffer (20 mM HEPES, pH 7.4, 500 mM NaCl, 10% glycerol, 5 mM imidazole, 2 mM β-mercaptoethanol, DNase1, and complete EDTA-free protease inhibitor). After disruption, the soluble fraction was separated by centrifugation and loaded onto a prepacked Ni-NTA column, equilibrated in the lysis buffer. The protein was eluted with an imidazole gradient. Prior to TEV protease cleavage, eluent fractions were dialyzed against cleavage buffer (50 mM Tris, pH 8.0, 150 mM NaCl, 2.5 mM CaCl_2_). After complete digestion, Ni-NTA affinity chromatography was performed again to remove undigested protein fractions. Size exclusion chromatography was then carried out using a Hi-Load 16/600 Superdex75 GL column to obtain a pure, monodispersed sample. After concentration, the protein was stored at -80 °C in storage buffer (20 mM Tris, pH 8.0, 150 mM NaCl, 5% glycerol, 5 mM dithiothreitol).

### Crystallization and structure determination

The MKP5-CD variant Y435W, at a concentration of 6-8 mg/ml, underwent co-crystallization with the Cmpd 1^17^ using the sitting-drop method. The protein was incubated overnight with Cmpd 1 (5 mM). Drops of 300 nl protein mixed with 200 nl reservoir solution were set using the Formulatrix NT8 drop setter in a 96-well plate. Crystals of the Y435W in complex with Cmpd 1 were grown in a buffer of 0.1 mM Tris, 300 mM NaCl, pH 8.5. X-ray crystallographic data sets were collected at Brookhaven Lab National Synchrotron Light Source II, at the AMX and FMX beamlines. Molecular replacement, employing either Molrep in the CCP4 software package or phaser^34^, was carried out using the MKP5-CD structure in complex with Cmpd 1 (PDB: 6MC1) as the search model. The 2|Fo| - |Fc| and |Fo| - |Fc| Fourier maps were analyzed, revealing no electron density for any compounds other than Cmpd 1. Energy-minimized coordinates and Crystal Information Files (CIF) for Cmpd 1 were generated using Jligand^35^. Structure refinement utilized Phenix.refine^36^ and REFMAC5^37^, with several rounds of refinement cycles and model building in Coot^38^ to achieve the final structure. Stereochemistry assessment with MOLPROBITY^39^ confirmed good quality in the crystal structures. Data collection, scaling, and refinement statistics are summarized in (Supplementary Table 1). The structural coordinates of the complex have been deposited in the RCSB Protein Data Bank PDB (9BPN: apo-Y435W, 9BU4: Cmpd 1-Y435W).

### NMR spectroscopy

MKP5-CD samples for NMR were expressed in M9 minimal media with ^15^NH_4_Cl and ^13^C_6_H_12_O_6_ (Cambridge Isotope Laboratories) as the sole nitrogen and carbon sources. The cells were cultured at 37 °C to an OD_600_ of 0.8 – 1.0, induced with 1 mM IPTG, and then incubated at 20 °C for an additional 16 – 18 hours. Isotopically labeled MKP5-CD samples were purified by the protocol described above. The samples were dialyzed into a buffer containing 40 mM sodium phosphate, 2 mM DTT, and 1 mM EDTA at pH 7.4. Samples were concentrated to 0.1 – 0.5 mM and 7.5% D_2_O was added prior to NMR experiments. For studies of Cmpd 1 binding, 0.1 mM MKP5-CD was incubated with 0.12 mM Cmpd 1 (dissolved in 100% DMSO) for 1 hour at room temperature prior to data collection and compared to spectra of 0.1 mM MKP5-CD (apo). All NMR experiments were collected on a Bruker Avance NEO 600 MHz spectrometer at 25°C equipped with pulsed field gradients and a triple resonance cryoprobe. NMR spectra were processed with NMRPipe^40^ and analyzed in Sparky^41^. The backbone resonance assignments were completed using a TROSY HSQC and the following TROSY triple resonance experiments: HNCA, HN(CO)CA, HN(CA)CB, HN(COCA)CB, HN(CA)CO, and HNCO^42, 43^. Three-dimensional correlations and resonance assignments were determined in CARA^44^. 37.3% of the MKP5-CD sequences remain unassigned due to line broadening that precludes reliable visualization of these resonances. ^1^H^15^N combined chemical shift perturbations (Δδ) were determined from ^1^H^15^N TROSY HSQC spectra by 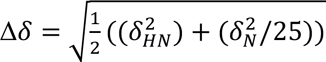, where δ_HN_ and δ_N_ = |*δ_WT_* – *δ_variant_*|. Significant chemical shift perturbations were determined as values 1.5σ above the 10% trimmed mean of all data sets.

### CD Spectroscopy

MKP5-CD samples (10 μM) were dialyzed into a buffer containing 40 mM sodium phosphate and 1 mM EDTA at pH 7.4. Thermal unfolding circular dichroism (CD) experiments were collected on a JASCO J-815 spectropolarimeter equipped with a variable temperature Peltier device using a 2 mm quartz cuvette. Denaturation curves were recorded at 223 nm over a temperature range of 20 – 90 °C. *T_m_* values were determined via nonlinear curve fitting in GraphPad Prism.

### Size Exclusion Chromatography

To assess the oligomeric state of MKP5 in complex with Cmpd 1, 400 μL of 65 μM WT MKP5 was run on an Enrich SEC 650 size exclusion column (Bio-Rad) at 1 mL/min in a buffer containing 40 mM sodium phosphate, 2 mM DTT, and 1 mM EDTA at pH 7.4 (consistent with NMR studies). The samples included (1) an apo WT MKP5 sample and (2) a WT MKP5 sample incubated with 1.2 molar excess Cmpd 1 (78 μM) for 1 hour at room temperature prior to chromatography.

### Molecular Dynamics simulations

The structural models for the apo and Cmpd 1-bound protein of MKP5 were based on the crystal structure of MKP5 with (PDB: 6MC1)^17^ and without Cmpd 1 (PDB: 1ZZW)^20^. MD simulations were performed using the AMBER-ff19SB, and General Amber (GAFF) force field for the protein and ligand, respectively, as included in the Amber22 software package^45, 46^. We simulated the dynamics of eight distinct states of MKP5, including WT and three variants (Y435W, Y435A, Y435S). Additionally, WT and Y435W were simulated in the presence of Cmpd 1. MD simulations over 1 µs were carried out for each system. AmberTools22 were employed to define standard protonation states and complete hydration of the models using TIP3 water solvent molecules to ensure density values ≥0.9 mol Å^−3^ ^46^.

MD simulations were prepared as follows: first, the solvent was minimized by restraining all atoms but water and ions at the positions defined by the PDB models. Structural constraints were then released, and the solvated structures were then gradually heated to 303 K, performing MD simulations (of at least 10 ns) in the canonical NVT ensemble using Langevin dynamics. Unconstrained MD simulations were carried out for 90 ns, resulting in a total pre-equilibration simulation time of about 100 ns. The pre-equilibrated model systems were then simulated in the NPT ensemble at 300 K and 1 atm, using Langevin dynamics for 1µs. All simulations were performed using periodic boundary conditions with a switching cutoff distance of 12 Å. Three independent replicas were run for WT and each variant in the unbound forms and WT and Y435W bound forms for a total of 18 µs simulations analyzed in this study.

The MD trajectories were analyzed using the MDiGest^47^ and MDAnalysis^48^ Python packages after sampling one snapshot per 0.1 ns (i.e., a total of 10000 frames for each model system). The root-mean-square deviation (RMSD) analysis and non-covalent interaction analyses were performed as implemented in MDiGest^47^. For the latter, 1 frame per ns and the default angular and distance settings were used to estimate the percentage of frames in which each interaction was present^47^.

The traffic maps in Figure 3i use a new implementation based on a modified Dijkstra algorithm^49^ which finds the ten highest correlated paths connection between each pair of residues in the protein. To do so, the protein is represented as a graph where each amino acid corresponds to a node, and the edges connecting them are proportional to the computed generalized correlation values of backbone dihedral fluctuations obtained from the MD simulations. Hence, each residue is represented as a vector of four elements corresponding to the sine and cosine angles projections of phi and psi backbone dihedral angles computed at each step of the trajectory. The resulting time series is used to compute the Mutual-information based generalized correlation matrix, as implemented in MDigest^47^. The resulting matrix is sparsified to obtain the network for the ten highest correlated-path-computation. Only edges corresponding to pairs of residues whose Cα distances are at less than 5 Å during the whole MD simulation time are kept. Finally, the –log transformation is applied to the resulting matrix, such that high correlation edges map to short edge distances, whereas less correlated residue pairs (values closer to 0) correspond to long edge distances. The resulting graph with nodes and edges based on proximity and correlation is used as input for the traffic analysis, resulting in the computation of the ten shortest and suboptimal paths between each pair of nodes. The total edge traffic *e_t_* for each edge (between nodes *i*, *j*) is computed as 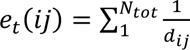, where *d_ij_* is the length of the corresponding path and the summation runs on the number of times each edge is sampled. Finally, the edge-traffic values are computed separately from the apo trajectories of each system WT, Y435W, Y435S, and Y435A, and normalized with respect to the maximum value over all variants. The resulting edge-traffic values are represented on the protein structure. Only the edges *e_t_* (*ij*) greater than 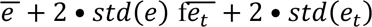 for each system are displayed.

### Generation of MKP5 mutant expression vectors and cell transfections

Generation of MKP5 mutant variants, MKP5-Y535S, MKP5-Y435A and MKP5-Y435W have all been described previously^17^. Cell-based assays for MAPK binding experiments utilized COS-7 cells grown at 37°C at 5% CO_2_. Cells were maintained in Dulbecco’s modified eagle’s medium (DMEM) (cat# 11965092) supplemented with 10% fetal bovine serum (FBS), 1% sodium pyruvate (Gibco cat# 11360070), and 1% penicillin-streptomycin (Gibco cat# 15140122). COS-7 cells were seeded at a density of 8.5x10^5^ in 100 mm dishes and maintained in DMEM supplemented with 10% FBS for 24 h before transfection. Cells were co-transfected with 6.8 µg DNA of either pEBG vector (expressing GST alone) or full-length pEBG-GST-MKP5 (WT or variants), and 3.4 µg of either pcDNA3-HA-p38α MAPK or pEGFP-JNK1β using FuGENE transfection reagent as per manufacturer’s instructions. Cells were transfected for 48 h in 10% FBS DMEM before harvest.

### Immunoblotting and immunoprecipitation

Cells were lysed on ice using Pierce^TM^ IP lysis buffer (cat# 87787). Lysates were affinity purified using GSH-Sepharose beads. Affinity purified complexes containing full-length MKP5 (WT or variants) were separated by boiling and complexed HA-p38α MAPK or pEGFP-JNK1β proteins detected by immunoblotting using either anti-HA (3F10, Roche cat# 121518167001) or anti-GFP (Cell Signaling Technologies cat# 2955) antibodies. GST input controls were detected using anti-GST antibodies. For the assessment of MAPK phosphorylation, cell lysates were clarified by centrifugation, and proteins were resolved by SDS-PAGE and transferred to nitrocellulose membranes by semi-dry transfer for 1 h followed by immunoblot analysis for detection of endogenous phosphorylated p38 MAPK, JNK1/2, and ERK1/2 using anti-phospho-p38 MAPK (Cell Signaling Cat# 9215), anti-phospho-JNK1/2 (Cell Signaling cat#4668) and anti-phospho-ERK1/2 (Cell Signaling cat# 9101) antibodies. Total levels of MAPKs were assessed by immunoblotting of re-probed membranes using anti-p38α MAPK (Santa Cruz cat# 81621), anti-JNK1/2 (Santa Cruz cat# 3708), and anti-ERK1/2 (Cell Signal cat# 9107) antibodies.

### Statistical analyses

No statistical methods were used to predetermine sample size. The number of samples used in each experiment is shown. All cell-based experiments were performed at least three times independently. Statistical analysis and graphing were performed using GraphPad Prism 9.4.1 software. We did not estimate variations in the data. The variances are similar between the groups that are being statistically compared. All data represent the means ± standard errors of the means (SEM). For p-value determinations, we used one-way or two-way ANOVA with multiple comparisons.

## ACKNOWLEDGMENTS

This work was partially supported by NIH grant R01 GM144451 to G.P.L. A.M.B and E.L. were supported by R01 HL158876, A.M.B. was also partially supported by R01 AR080152.

## AUTHOR CONTRIBUTIONS

ES: expressed and purified MKP5-CD proteins, performed NMR and CD experiments and associated data analysis; FM: performed MD simulations and associated data analysis; RM: expressed and purified MKP5-CD proteins, performed X-ray crystallography and associated data analysis; SS: performed MKP5 cellular experiments and associated data analysis; EJL: conceived the study, supervised X-ray crystallography; VSB: conceived the study, supervised MD simulations; AMB: conceived the study, supervised cellular experiments; GPL: conceived the study, supervised NMR spectroscopy. The manuscript was written through contributions of all authors.

## DATA AVAILABILITY

All data needed to evaluate the conclusions in the paper are present in the paper and/or the Supplementary Materials.

---

**Supplementary Figure 1.**
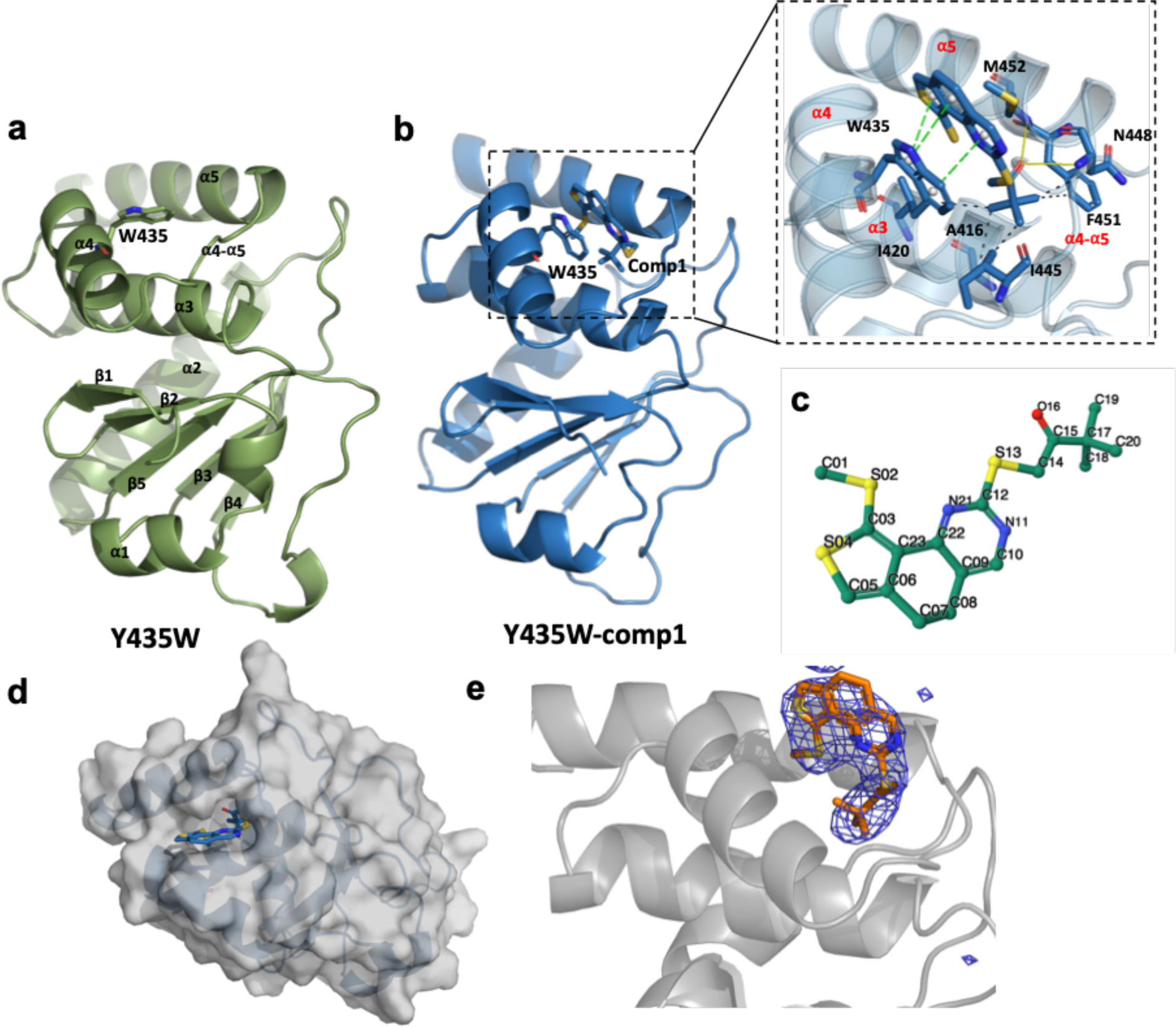
Structures of Y435W mutants with and without Cmpd 1 bound. (**a)** The apo Y435W structure (green), with the mutated tryptophan residue within the allosteric pocket. **(b)** The Cmpd 1-bound Y435W structure (blue) highlights the allosteric site with the bound ligand (Cmpd 1) within the allosteric pocket. An enlarged view of the allosteric pocket to the right highlights the interacting residues. **(c)** Stick representation of Cmpd 1. (**d**) Surface representation of Y435W showing Cmpd 1 (blue sticks) bound in the allosteric pocket. (**e**). 2Fo-Fc map surrounding Cmpd 1.

**Supplementary Figure 2.**
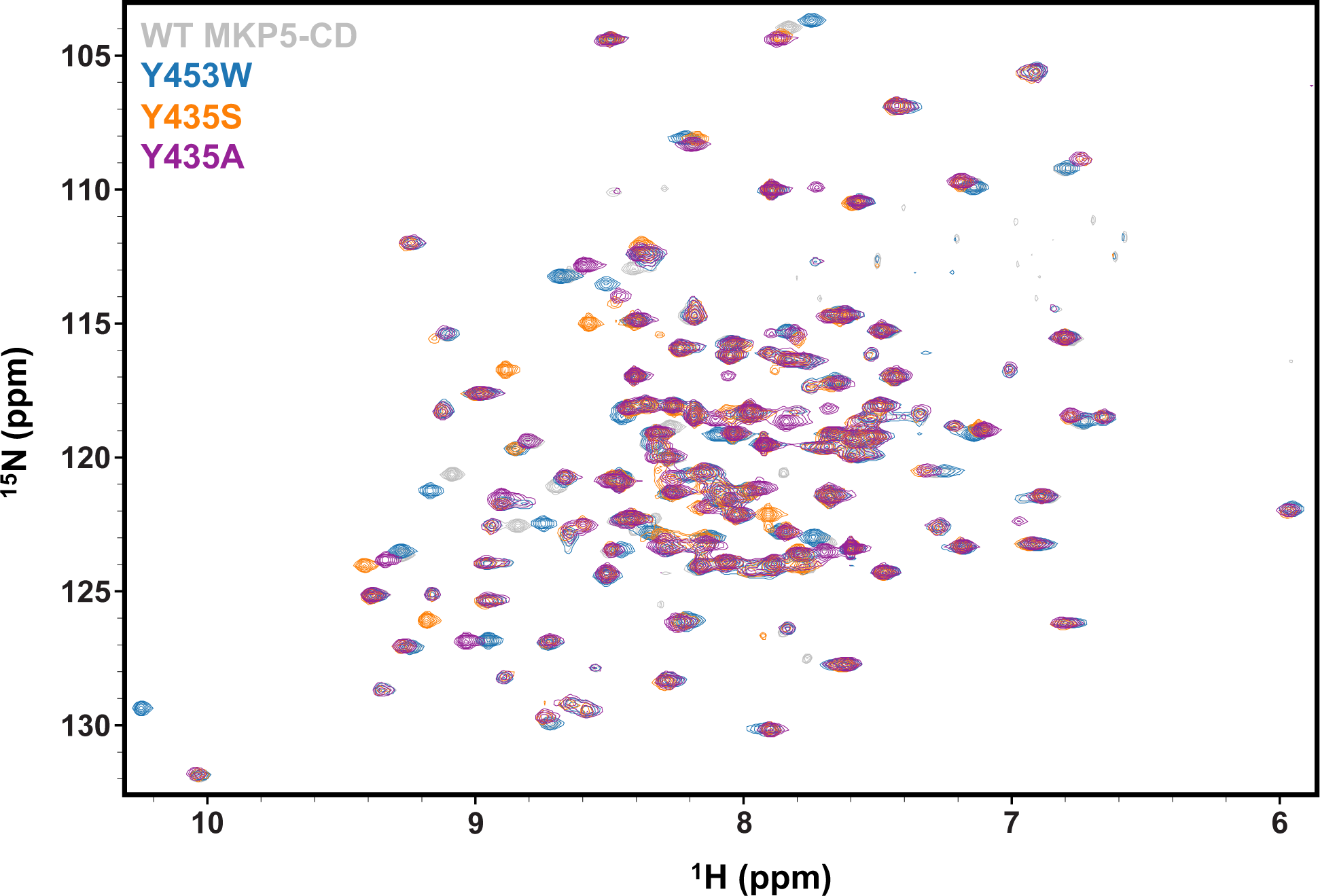
^1^H^15^N HSQC spectral overlay of WT MKP5-CD and the Y435 variants. WT MKP5-CD is shown in gray, Y435W in blue, Y435S in orange, and Y435A in purple.

**Supplementary Figure 3.**
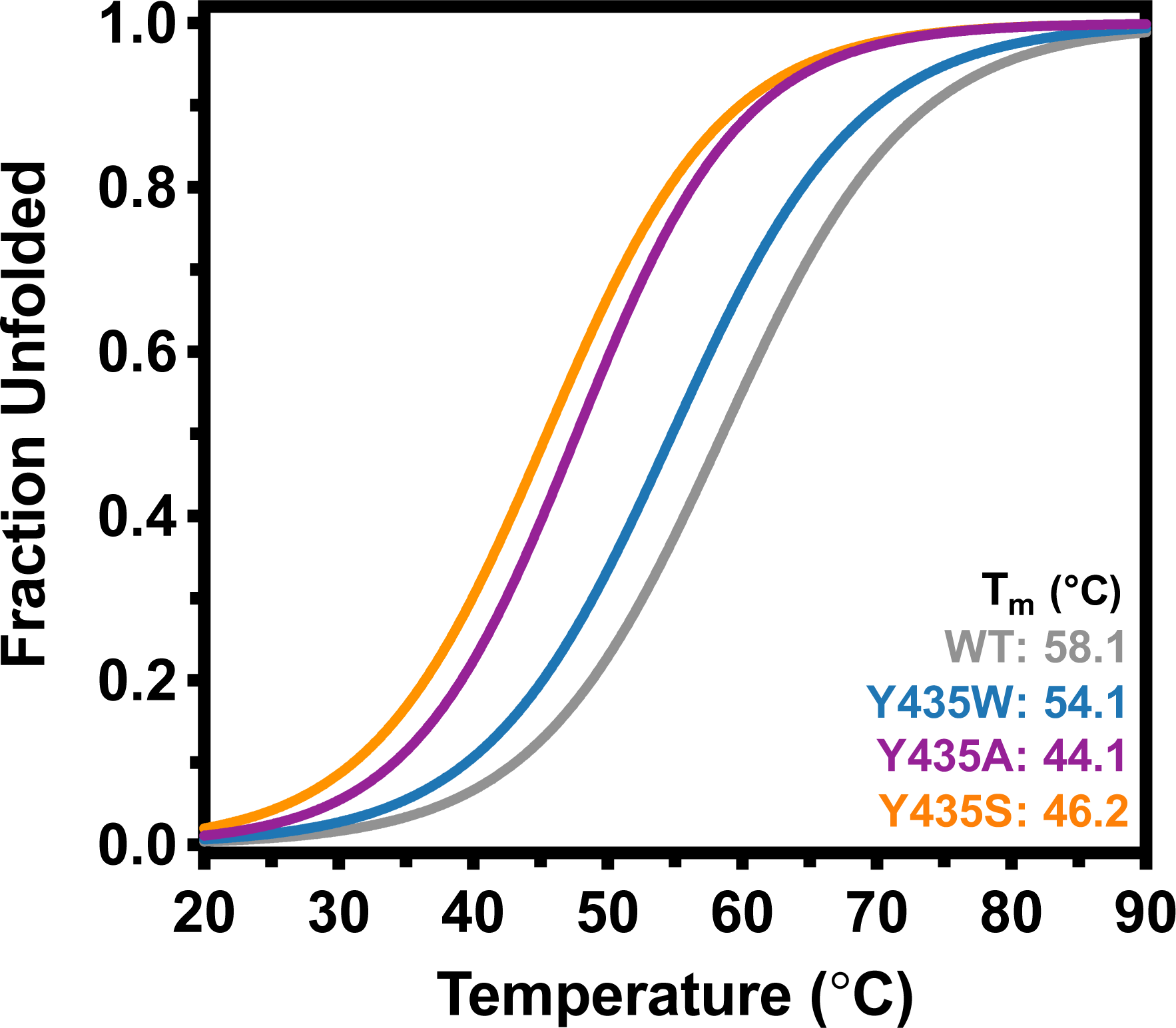
Thermal stability of WT MKP5-CD and the Y435 variants. Thermal unfolding curves derived from CD spectroscopy are shown for WT MKP5-CD (gray), Y435W (blue), Y435A (purple), and Y435S (orange). *T*m values were determined via nonlinear curve fitting in GraphPad Prism.

**Supplementary Figure 4.**
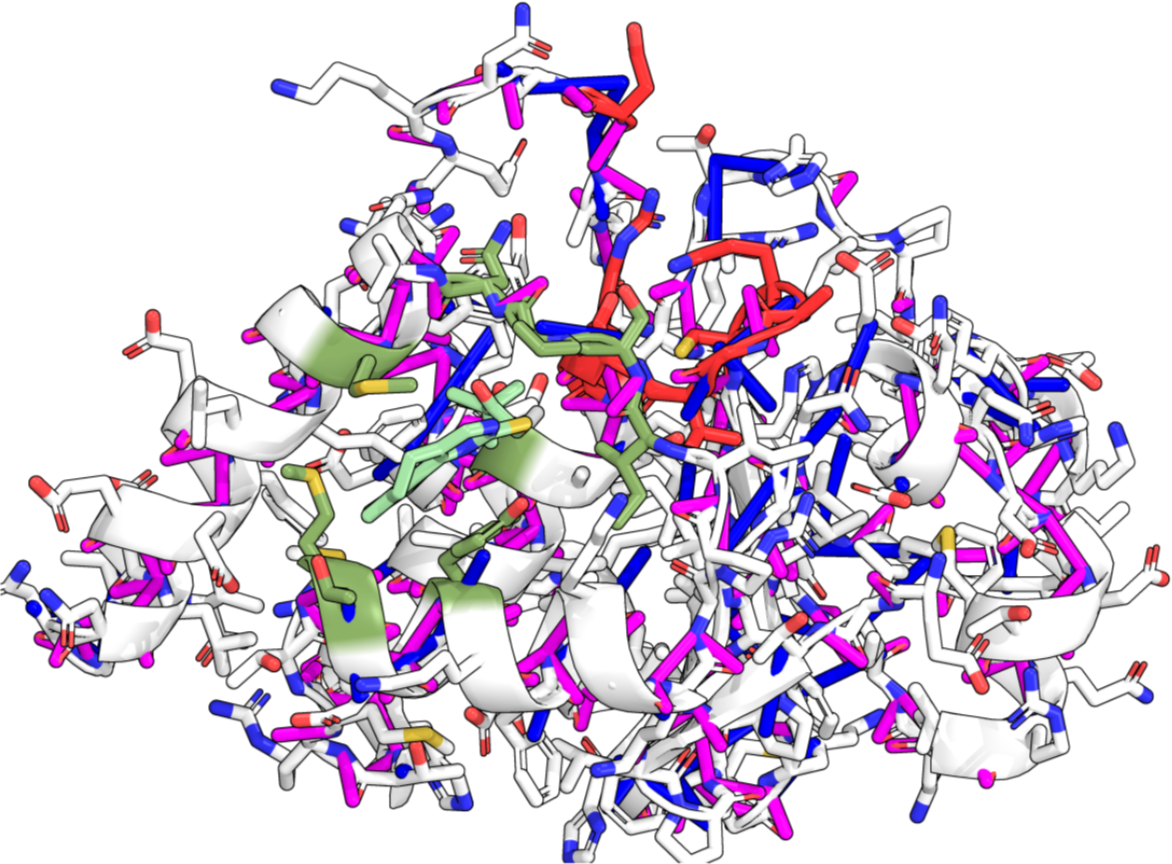
Unperturbed interaction network computed from 3 µs using WT apo trajectory (one frame per 10 ns). Polar (magenta) and hydrophobic (blue) interactions are shown in solid lines. Interactions with persistency exceeding 95% of total frames used are shown. The active site is shown in red sticks, while the allosteric site is shown in green sticks. Cmpd 1 is bound in the pocket and shown in light-green sticks.

**Supplementary Figure 5.**
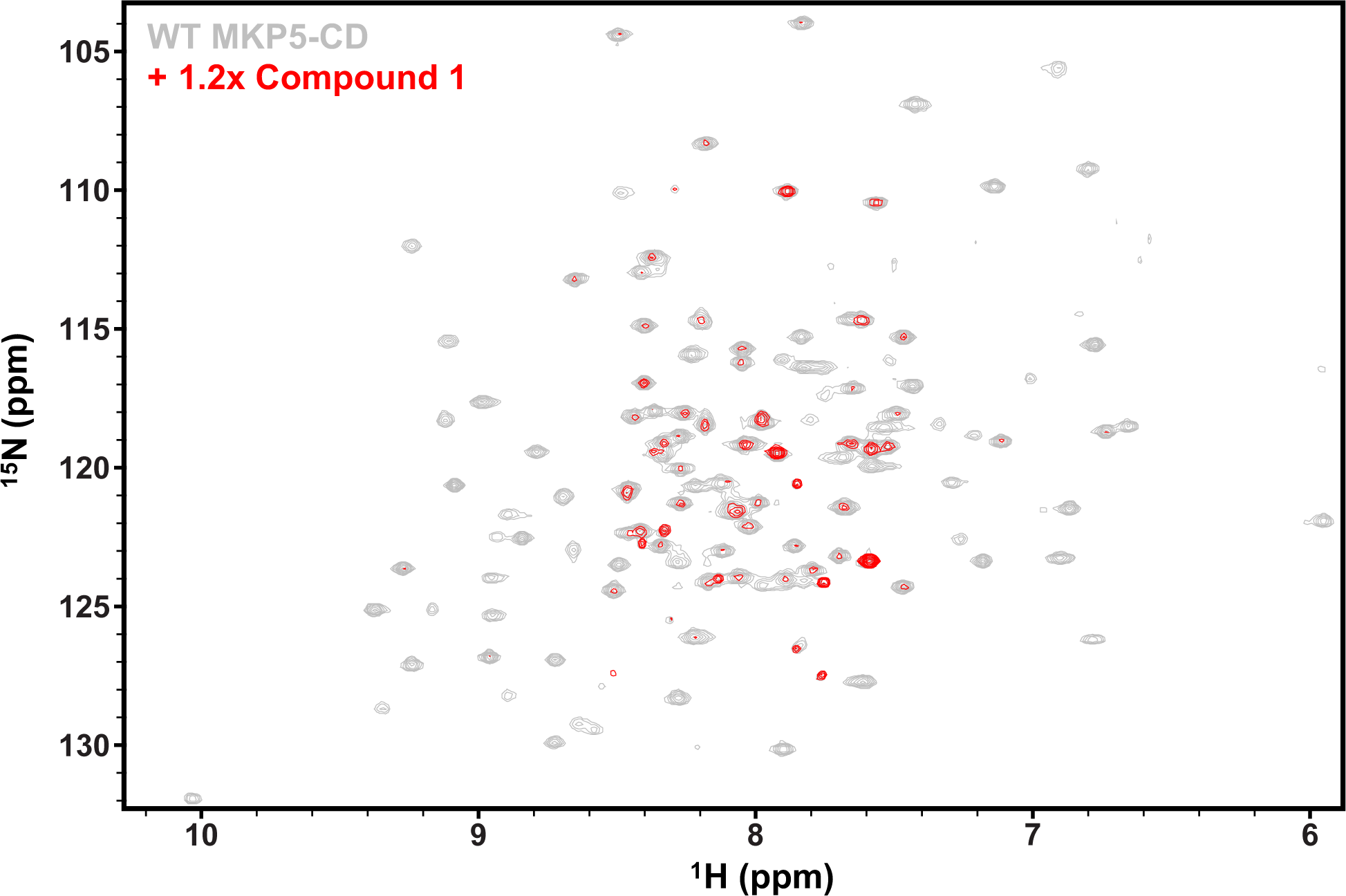
Effects of Cmpd 1 binding on the WT MKP5-CD structure via NMR. ^1^H^15^N HSQC spectra of apo WT MKP5-CD and Cmpd 1-bound WT MKP5-CD are overlaid in gray and red, respectively. Cmpd 1 (120 μM) was added in excess of WT MKP5-CD (100 μM).

**Supplementary Figure 6.**
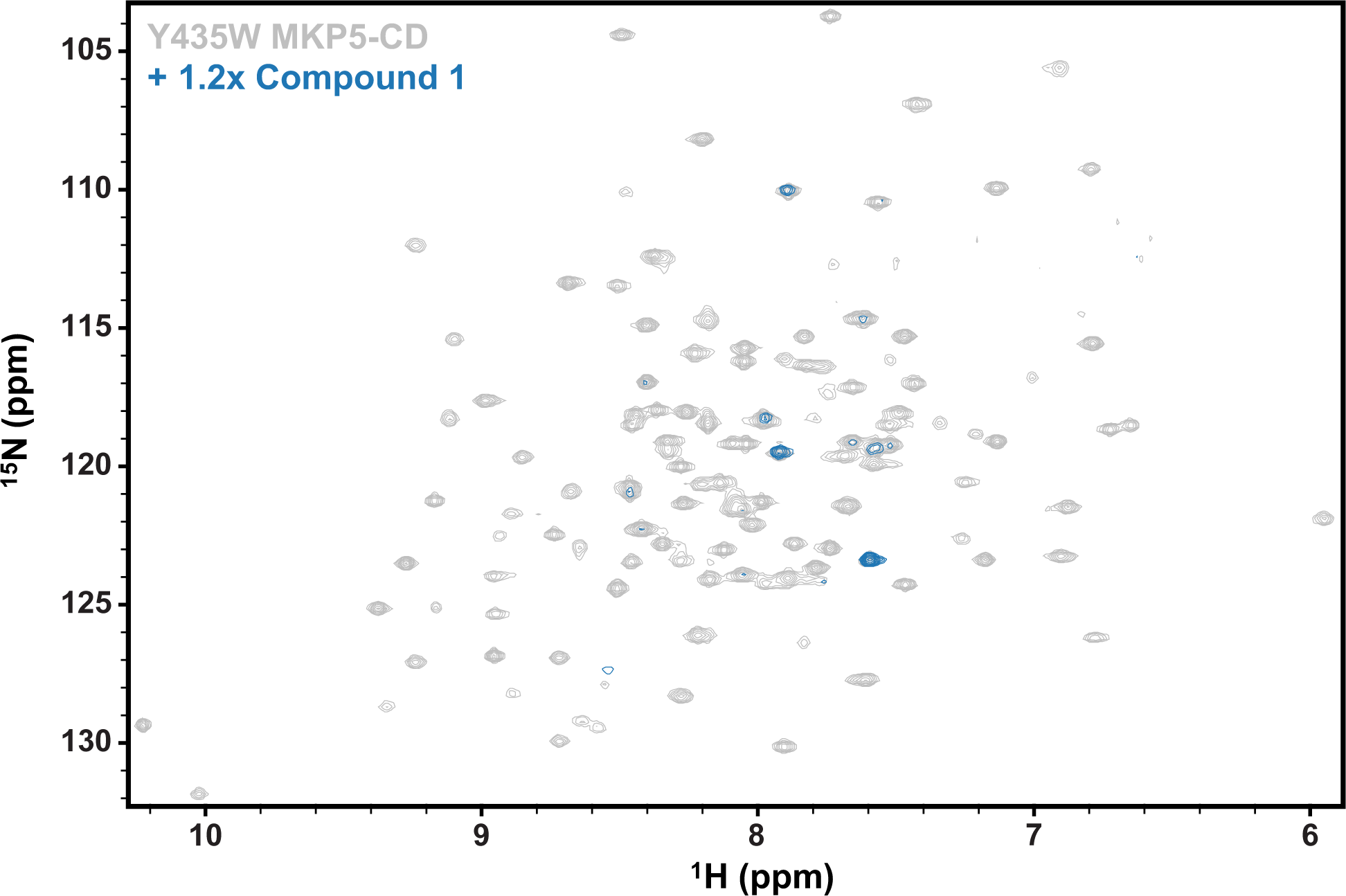
Effects of Cmpd 1 binding on the Y435W MKP5-CD structure via NMR. ^1^H^15^N HSQC spectra of apo Y435W MKP5-CD and Cmpd 1-bound Y435W MKP5-CD are overlaid in gray and blue, respectively. Cmpd 1 (120 μM) was added in excess of Y435W MKP5-CD (100 μM).

**Supplementary Figure 7.**
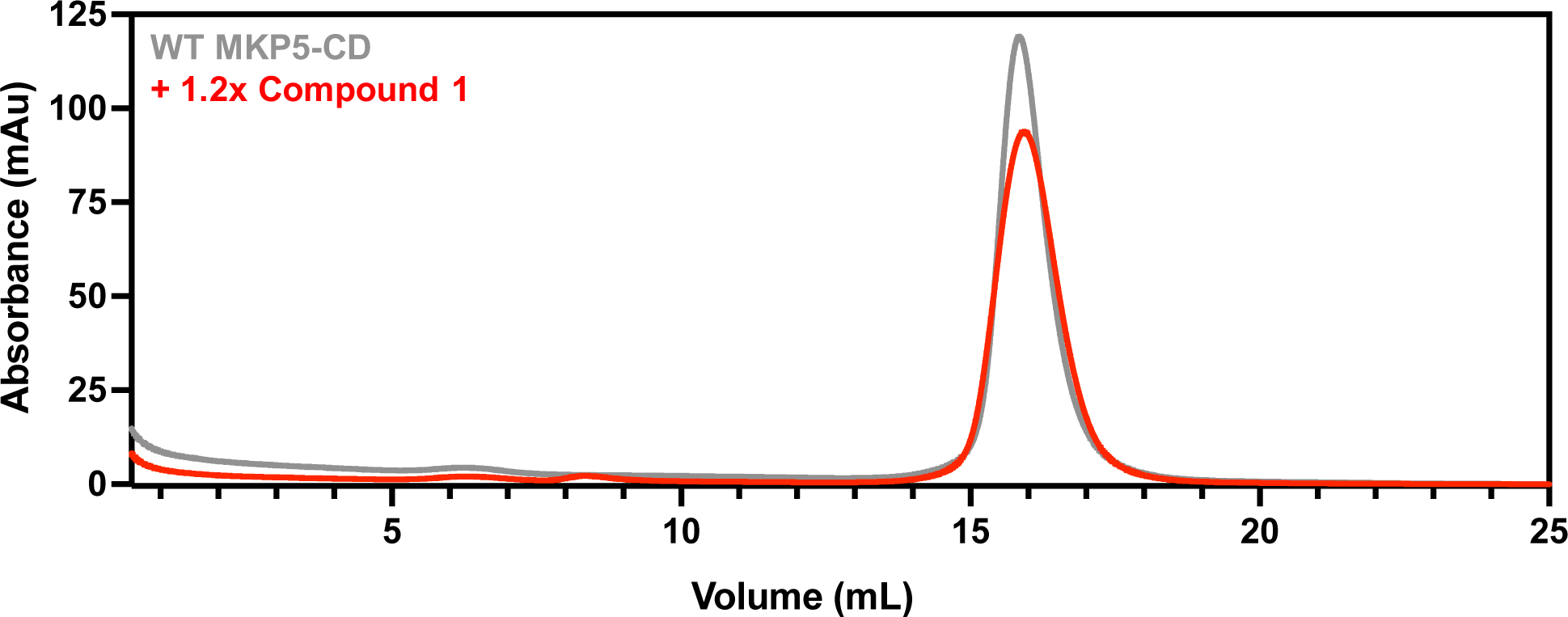
The binding of Cmpd 1 does not alter the oligomeric state of MKP5-CD. The size exclusion chromatogram of WT MKP5-CD (gray) is overlaid with that of WT MKP5-CD in the presence of 1.2x Cmpd 1 (red). Data were collected on a ENrich SEC 650 size exclusion column (Bio-Rad).

**Supplementary Figure 8.**
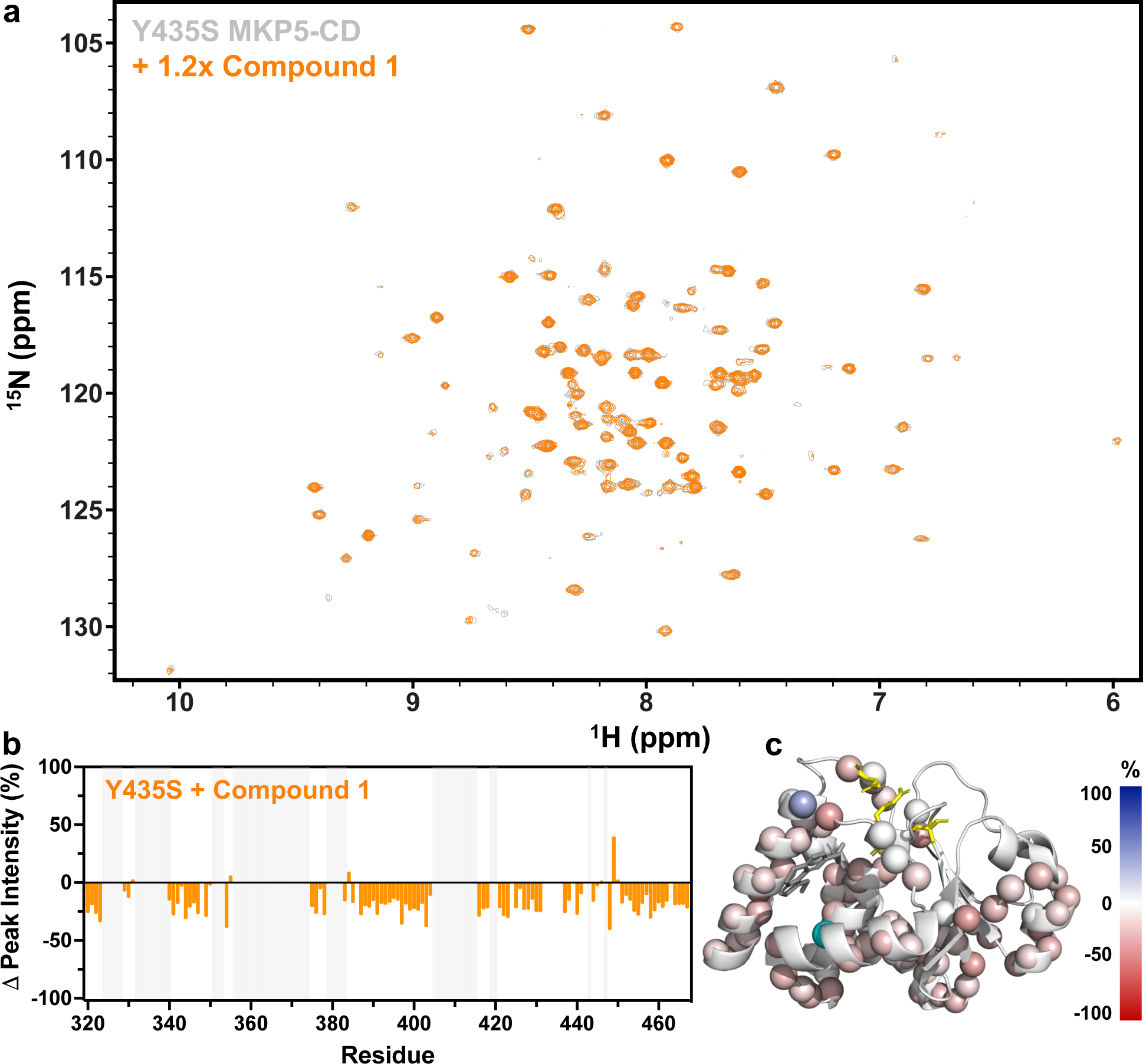
Effects of Cmpd 1 binding on the Y435S MKP5-CD structure via NMR. **(a)** ^1^H^15^N HSQC spectra of apo Y435S MKP5-CD and Cmpd 1-bound Y435S MKP5-CD are overlaid in gray and orange, respectively. Cmpd 1 (120 μM) was added in excess of Y435S MKP5-CD (100 μM). **(b)** Changes in ^1^H^15^N HSQC resonance (peak) intensity when Cmpd 1 is added to Y435S MKP5, calculated relative to apo Y435S MKP5. Light gray vertical lines on all plots indicate residues that are unassigned. **(c)** Δ peak intensity is plotted onto the MKP5 structure (PDB: 6MC1), with blue and red spheres representing decreases and increases in resonance intensity, respectively, and the color gradient denoting the magnitude of the change in resonance intensity in the presence of Cmpd 1 relative to apo MKP5. Residue 435 is represented by a cyan sphere, and the catalytic side-chains are highlighted in yellow. Cmpd 1 is modeled into the allosteric pocket for reference.

**Supplementary Figure 9.**
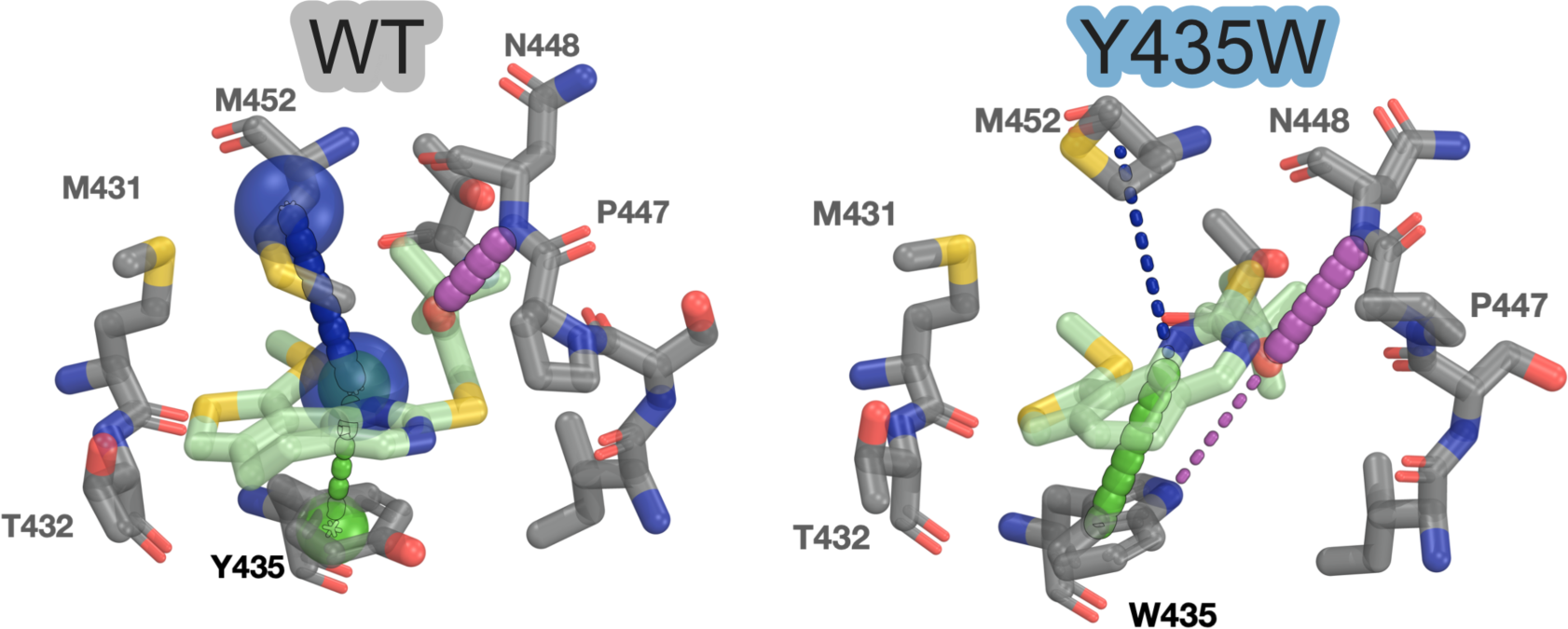
Binding pose interactions averages from 3 µs trajectories of WT and Y435W. Hydrophobic interactions are shown in blue, polar interactions in magenta, and pi-stacking interactions in green.

**Supplementary Figure 10.**
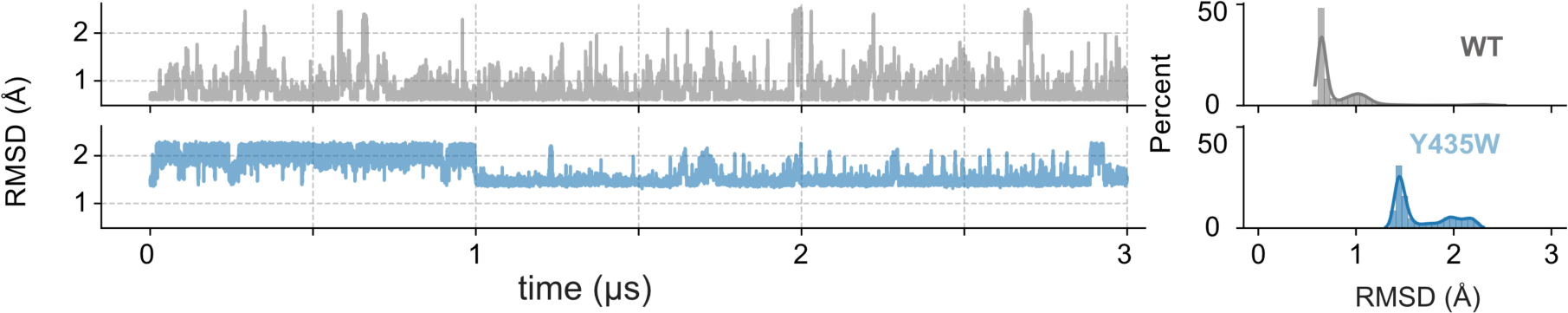
Effects of Cmpd 1 binding to Y435W MKP5. Root mean square deviations (RMSD) calculated along the trajectories of Cmpd-1 heavy atoms in WT and Y435W states are shown in the plots on the left, and corresponding density distributions, shown as percentages, are shown in the plots on the right.

**Supplementary Figure 11.**
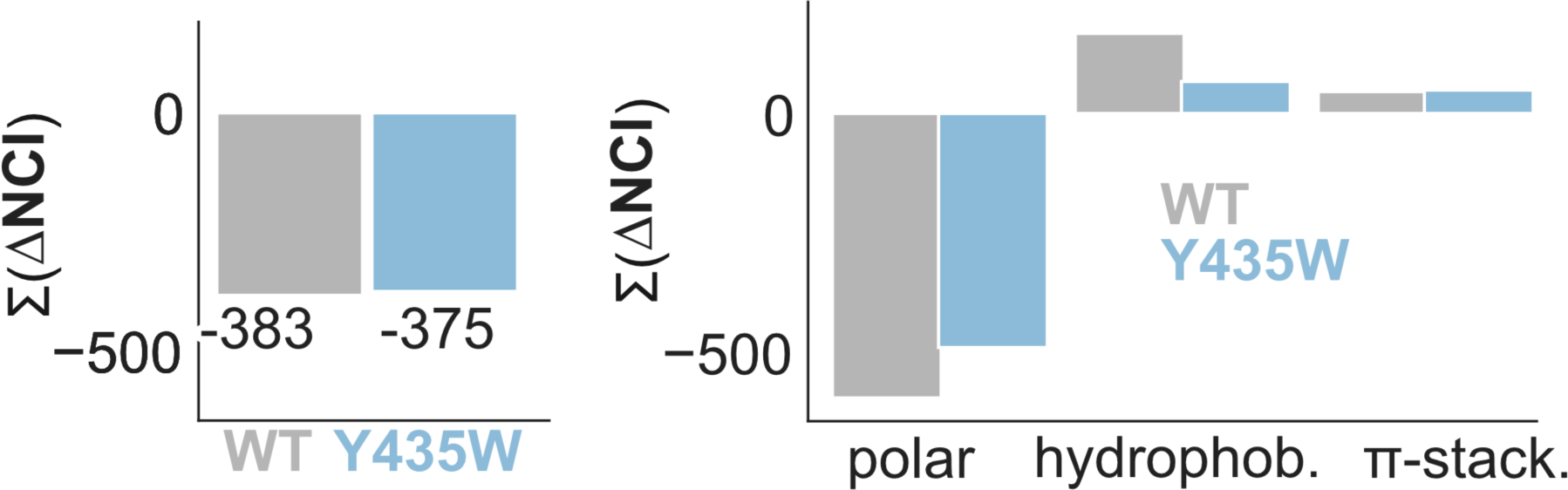
Cumulative persistence of non-covalent interaction (Σ(ΔNCI)) changes in WT and Y435W upon binding of Cmpd-1. The plot on the right shows ΔNCI changes decomposed into polar, hydrophobic and pi-stacking components.

**Supplementary Figure 12.**
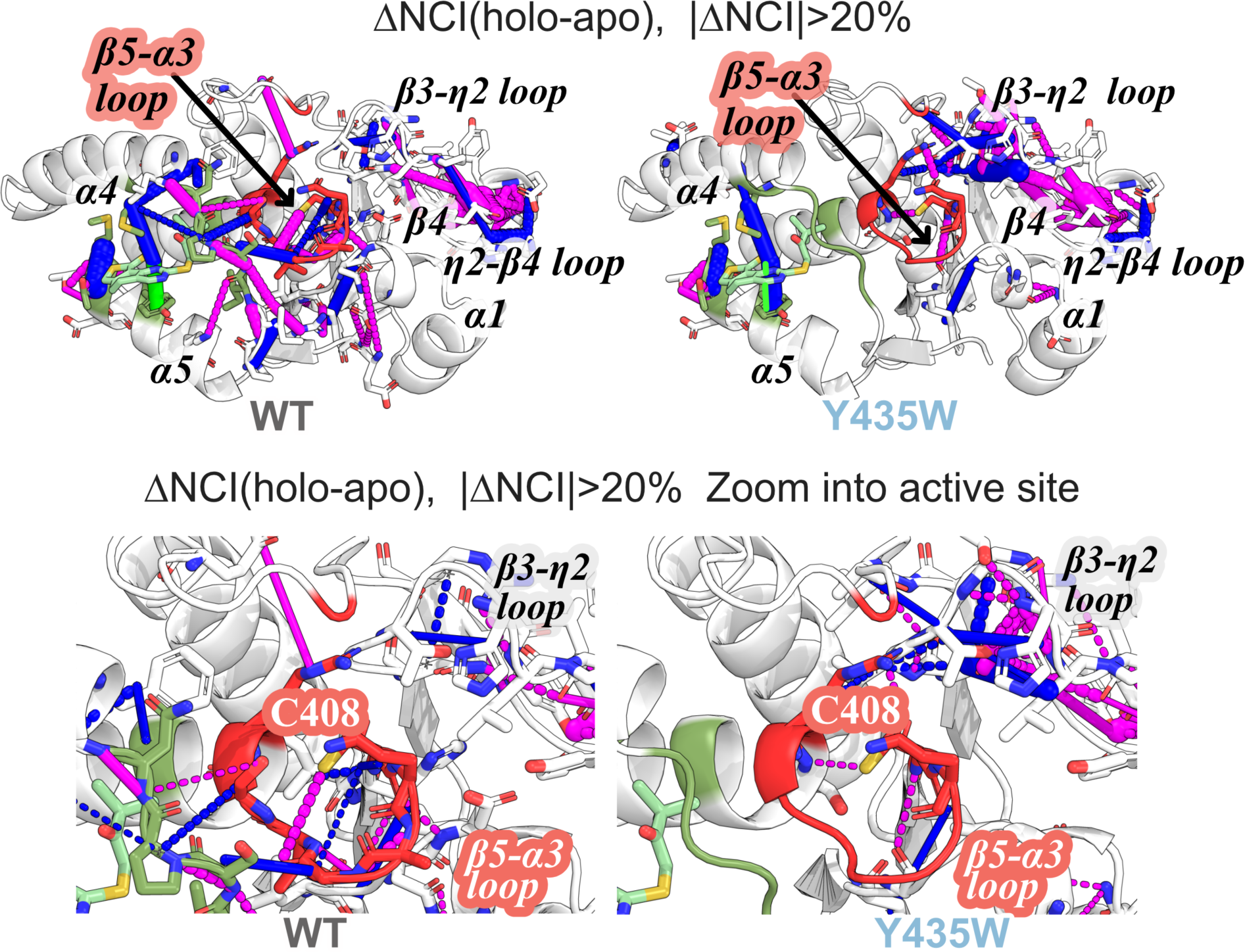
Reorganization of MKP5 non-covalent interactions upon allosteric site binding. **(top row)** Changes in non-covalent interactions (ΔNCI) sampled in WT and Y435W holo (Cmpd-1-bound) and apo states that change by more than 20% between holo and apo (unbound) states. **(bottom row)** Zoomed-in view of the active site regions. Mutant-specific differences are most abundant in the active and allosteric-site regions. Hydrophobic contacts are shown in blue, while polar contacts (hydrogen bonds and salt-bridges) are in magenta. Π-stacking interactions are shown in green. All non-covalent interactions are computed using 1 frame every ns over 3 µs simulated trajectories for each apo and holo states of WT and Y435W.

**Supplementary Table 1.** Data collection and refinement statistics for the apo-Y435W and Y435W-Cmpd 1 complex structures.

|  | **Apo-Y435W (9BPN)** | **Y435W-Cmpd 1(9BU4)** |
| --- | --- | --- |
| **Data collection** |  |  |
| Space group | C 1 2 1 | P 1 |
| Cell dimensions |  |  |
| *a*, *b*, *c* (Å) | 97.862 44.744 87.326 | 55.706 55.801 275.91 |
| $\alpha$, $\beta$, $\gamma$ (°) | 90 109.045 90 | 90.356 93.823 106.876 |
| Resolution (Å) | 27.52 - 2.40 (2.486 - 2.4) | 137.60 - 2.90 (2.97 - 2.90) |
| CC<sub>1/2</sub> | 0.995 (0.89) | 0.990 (0.247) |
| *I*/σ*I* | 10.57 (2.94) | 5.6 (0.8) |
| Completeness (%) | 99.80 (99.86) | 98.55 (98.4) |
| Redundancy | 6.8 (6.7) | 3.8 (3.9) |
| **Refinement** |  |  |
| Resolution | 28.00 - 2.40 (2.59 - 2.40) | 42.09 - 2.90 (3.004 - 2.90) |
| No. Reflections | 14234 (1421) | 69262 (6846) |
| *R*<sub>work</sub>/*R*<sub>free</sub> | 0.2364/0.2940 (0.2984/0.3453) | 0.2150/0.2662 (0.3528/0.3869) |
| No. non hydrogen atoms | 2292 | 14126 |
| Macromolecules | 2257 | 13829 |
| Ligand | - | 276 |
| B-factors (Å<sup>2</sup>) | 52.62 | 79.67 |
| Protein | 52.76 | 79.69 |
| Ligand | - | 80.83 |
| R.m.s. deviations |  |  |
| Bond lengths (Å) | 0.014 | 0.008 |
| Bond angles (°) | 1.42 | 0.96 |

Statistics for the highest-resolution shell are shown in parentheses.

**Supplementary Table 2.** Non-covalent interactions (NCI) persist for interactions changed by more than 15% in apo Y435W, Y435S, and Y435A compared to WT. hbond; hydrogen bond, hphob; polar interactions.

| donor | acceptor | WTapo | Y435W apo | Y435A apo | Y435S apo | type |
| --- | --- | --- | --- | --- | --- | --- |
| A375-N | T355-N | 43.03 | 26.59 | 31.27 | 32.50 | hbond |
| A410-N | G411-N | 43.44 | 26.39 | 20.13 | 22.50 | hbond |
| A435-OG | T432-N | NaN | NaN | NaN | 28.37 | hbond |
| A435-OG | T432-O | NaN | NaN | NaN | 15.33 | hbond |
| R372 | N353 | 68.15 | 89.09 | 86.33 | 58.93 | hphob |
| R372-NH1 | N353-OD1 | 44.47 | 73.98 | 69.43 | 38.72 | hbond |
| R372-NH1 | H357-N | 20.78 | 36.09 | 35.52 | 19.58 | hbond |
| R372-NH1 | T355-OG1 | 32.14 | 71.09 | 66.72 | 37.03 | hbond |
| R372-NH1 | T356-N | 17.90 | 27.98 | 35.43 | 17.73 | hbond |
| R372-NH2 | P359-N | 14.85 | 31.23 | 22.56 | 11.25 | hbond |
| R414 | V354 | 60.00 | 75.68 | 62.20 | 51.90 | hphob |
| R414-NE | Q409-O | 27.07 | 14.98 | 9.43 | 11.27 | hbond |
| R414-NH2 | D377-O | 4.54 | 2.48 | 2.48 | 19.57 | hbond |
| R414-NH2 | Q409-O | 23.31 | 11.11 | 6.75 | 7.53 | hbond |
| R414-NH2 | Q409-OE1 | 20.59 | 42.27 | 41.05 | 26.90 | hbond |
| R414-NH2 | S378-N | 1.87 | 1.50 | 1.25 | 20.40 | hbond |
| R414-NH2 | S378-O | 2.58 | 1.90 | 1.58 | 22.40 | hbond |
| R428-N | H426-N | 65.36 | 66.73 | 67.47 | 32.37 | hbond |
| R428-N | H426-O | 44.10 | 43.18 | 45.40 | 20.70 | hbond |
| R428-N | L423-O | 51.18 | 49.42 | 52.77 | 24.47 | hbond |
| R428-N | K425-O | 30.46 | 30.17 | 33.07 | 15.43 | hbond |
| R428-N | M424-O | 45.34 | 47.68 | 51.00 | 23.07 | hbond |
| R428-N | T427-N | 55.48 | 54.83 | 59.27 | 27.17 | hbond |
| R442-N | P443-N | 51.94 | 42.17 | 42.73 | 33.60 | hbond |
| R442-NE | I444-O | 13.05 | 34.31 | 32.30 | 27.63 | hbond |
| R442-NH1 | E321-O | 32.90 | 49.05 | 44.30 | 34.60 | hbond |
| R442-NH1 | I444-O | 9.67 | 2.41 | 3.15 | 32.55 | hbond |
| R442-NH1 | L331-O | 41.51 | 72.55 | 68.27 | 46.42 | hbond |
| R442-NH2 | N333-ND2 | 24.40 | 39.65 | 35.78 | 31.30 | hbond |
| N333-N | R442-NH2 | 20.49 | 38.53 | 29.97 | 29.13 | hbond |
| N353-ND2 | T355-OG1 | 10.56 | 32.13 | 30.50 | 12.62 | hbond |
| N382-N | Q381-NE2 | 45.99 | 67.70 | 59.70 | 71.93 | hbond |
| N382-N | K380-O | 24.20 | 35.94 | 32.63 | 39.33 | hbond |
| N448-N | L449-N | 23.02 | 44.95 | 51.07 | 48.83 | hbond |
| D341-N | Q338-O | NaN | NaN | NaN | 16.40 | hbond |
| D433-N | M431-O | 52.99 | 52.42 | 56.57 | 28.50 | hbond |
| D433-N | T430-O | 32.46 | 28.26 | 26.80 | 53.97 | hbond |
| D433-N | T432-OG1 | 39.28 | 39.87 | 43.40 | 20.10 | hbond |
| C408 | N333 | 0.40 | NaN | NaN | 31.30 | hphob |
| C408-SG | G411-O | 36.70 | NaN | NaN | NaN | hbond |
| C408-SG | S415-N | 17.39 | 33.43 | 26.07 | 21.47 | hbond |
| Q381 | S378 | 28.22 | 13.05 | 20.17 | 12.50 | hphob |
| Q381-NE2 | N382-N | 26.17 | 38.14 | 34.27 | 41.40 | hbond |
| Q381-NE2 | Q385-NE2 | 37.60 | 50.49 | 46.52 | 53.45 | hbond |
| Q381-NE2 | T376-O | 24.76 | 2.90 | 15.80 | 2.43 | hbond |
| Q409 | C408 | 59.36 | 78.34 | 87.77 | 86.70 | hphob |
| Q409-N | R414-NH2 | 25.62 | 15.98 | 14.30 | 8.43 | hbond |
| Q409-NE2 | R414-NH2 | 11.47 | 27.15 | 20.75 | 21.35 | hbond |
| Q454-NE2 | Q381-O | 40.42 | 48.75 | 61.43 | 48.62 | hbond |
| E334-N | Q409-O | NaN | 0.10 | 0.15 | 18.10 | hbond |
| G411 | C408 | 21.00 | NaN | NaN | NaN | hphob |
| G411-N | A410-N | 72.64 | 57.72 | 47.10 | 43.93 | hbond |
| G411-N | C408-O | 22.40 | 0.10 | NaN | NaN | hbond |
| 13 | V412-N | 67.02 | 89.02 | 91.57 | 90.60 | hbond |
| H357-N | L358-N | 79.30 | 95.46 | 91.80 | 77.93 | hbond |
| H357-N | T355-OG1 | 26.54 | 42.67 | 42.30 | 27.73 | hbond |
| H357-N | T356-OG1 | 47.43 | 73.17 | 67.53 | 52.20 | hbond |
| H407 | N353 | 0.30 | 0.25 | 7.40 | 22.80 | hphob |
| I444 | P443 | 38.30 | 72.81 | 68.47 | 62.00 | hphob |
| I444-N | I445-N | 41.74 | 3.24 | 8.57 | 14.63 | hbond |
| I445-N | P443-O | 62.64 | 33.03 | 36.10 | 36.60 | hbond |
| L331-N | F330-N | 53.22 | 64.39 | 53.83 | 36.20 | hbond |
| L340 | A337 | NaN | NaN | NaN | 24.00 | hphob |
| L340-N | Q338-N | NaN | NaN | NaN | 22.20 | hbond |
| L358 | H357 | 53.86 | 80.38 | 74.63 | 84.13 | hphob |
| L449-N | N448-N | 32.60 | 62.97 | 66.23 | 64.20 | hbond |
| L449-N | N448-OD1 | 54.11 | 77.35 | 74.23 | 77.50 | hbond |
| K436-N | A435-OG | NaN | NaN | NaN | 15.37 | hbond |
| K439-NZ | I445-O | 24.22 | 8.71 | 7.15 | 4.84 | hbond |
| K439-NZ | P443-O | 33.69 | 34.00 | 16.60 | 14.26 | hbond |
| K441 | I325 | 25.56 | 39.90 | 42.73 | 26.77 | hphob |
| M429-N | M424-O | 36.66 | 35.43 | 34.13 | 21.03 | hbond |
| M431-N | T430-OG1 | 64.14 | 66.80 | 66.07 | 39.37 | hbond |
| M431-N | T432-N | 46.27 | 43.14 | 39.87 | 76.70 | hbond |
| M452 | F451 | 39.05 | 18.12 | 13.10 | 7.23 | hphob |
| M452 | P447 | 20.43 | 0.90 | 0.60 | 0.60 | hphob |
| F451 | N448 | 59.19 | 86.32 | 84.70 | 87.17 | hphob |
| P443 | R442 | 45.00 | 59.86 | 60.97 | 67.20 | hphob |
| P447 | S413 | 29.00 | 3.10 | 0.15 | 0.50 | hphob |
| S413-OG | N448-N | 25.94 | 5.44 | 3.07 | 2.50 | hbond |
| S446 | G411 | 43.41 | 75.17 | 71.97 | 73.67 | hphob |
| S446-N | I444-O | 32.64 | 1.10 | 2.80 | 9.07 | hbond |
| S446-N | P447-N | 42.47 | 2.60 | 1.33 | 1.75 | hbond |
| T355 | V354 | 29.54 | 44.41 | 45.47 | 30.27 | hphob |
| T355-OG1 | N353-ND2 | 30.32 | 57.76 | 51.53 | 34.00 | hbond |
| T355-OG1 | Q409-NE2 | 19.50 | 2.80 | 28.40 | 5.90 | hbond |
| T355-OG1 | Q409-OE1 | 21.00 | 1.20 | 19.90 | 2.27 | hbond |
| T355-OG1 | L358-N | 17.10 | 35.44 | 32.77 | 26.30 | hbond |
| T427-N | R428-N | 91.04 | 90.77 | 96.33 | 44.57 | hbond |
| T427-N | M424-O | 16.92 | 14.88 | 12.77 | 33.27 | hbond |
| T427-N | Y422-O | 35.87 | 39.17 | 39.73 | 19.77 | hbond |
| T430-OG1 | D433-N | 21.91 | 16.85 | 14.80 | 63.60 | hbond |
| T430-OG1 | D433-OD2 | 10.54 | 9.51 | 8.43 | 26.00 | hbond |
| T430-OG1 | M431-N | 63.32 | 67.34 | 67.43 | 14.00 | hbond |
| T432-N | T430-OG1 | 46.34 | 46.14 | 41.47 | 62.77 | hbond |
| T467 | D461 | 25.38 | 25.79 | 20.93 | NaN | hphob |
| Y361 | L360 | 1.20 | NaN | 0.50 | 19.13 | hphob |
| Y361-N | L360-N | 93.57 | 97.56 | 94.40 | 78.43 | hbond |
| Y370-OH | N353-ND2 | 4.80 | NaN | NaN | 20.70 | hbond |
| Y370-OH | H362-N | 62.67 | 61.83 | 64.03 | 44.27 | hbond |
| Y370-OH | Y363-N | 76.88 | 79.85 | 80.87 | 55.70 | hbond |
| V354 | N353 | 61.79 | 88.46 | 83.73 | 75.63 | hphob |
| V412-N | N333-OD1 | 0.10 | 0.10 | 0.30 | 18.70 | hbond |
| V412-N | G411-N | 55.71 | 72.90 | 73.60 | 73.57 | hbond |
| V412-N | S413-N | 65.12 | 45.74 | 45.83 | 48.07 | hbond |

